# Ubiquitin ligase TRAIP plays an essential role during the S-phase of unperturbed cell cycle in the resolution of DNA replication – transcription conflicts

**DOI:** 10.1101/2022.03.23.485338

**Authors:** Shaun Scaramuzza, Martina Muste Sadurni, Divyasree Poovathumkadavil, Toyoaki Natsume, Patricia Rojas, Masato T. Kanemaki, Marco Saponaro, Agnieszka Gambus

**Author notes:** Research Center for Genome & Medical Sciences, Tokyo Metropolitan Institute of Medical Science, Tokyo, Japan (Toyoaki Natsume).

## Abstract

Cell division is the basis for the propagation of life and requires accurate duplication of all genetic information. DNA damage created during replication (replication stress) is a major cause of cancer, premature aging and a spectrum of other human disorders. Over the years, TRAIP E3 ubiquitin ligase has been shown to play a role in various cellular processes that govern genome integrity and faultless segregation. TRAIP is essential for cell viability, and mutations in TRAIP ubiquitin ligase activity lead to primordial dwarfism in patients. Here, for the first time, we have determined the mechanism of inhibition of cell proliferation in TRAIP-depleted cells. We have taken advantage of the auxin induced degron system to rapidly degrade TRAIP within cells and to dissect the importance of various functions of TRAIP in different stages of the cell cycle. We find that upon rapid TRAIP degradation, specifically in S-phase, cells cease to proliferate, arrest in G2 stage of the cell cycle and undergo senescence. Our findings reveal that TRAIP works in S-phase to prevent DNA damage at transcription start sites, caused by replication-transcription conflicts.

## Introduction

Cell proliferation is the basis for the propagation of life and requires accurate duplication of all genetic information before correct cell division. However, these processes encounter impediments that threaten their faultless execution and lead to genomic instability. Moreover, problems encountered during DNA replication (replication stress) often result in underreplicated DNA that needs to be resolved during mitosis. Cells have developed a number of means to respond to these challenges, such as replication-coupled DNA repair pathways and S-phase checkpoint responses that protect the stability of replication forks, inhibit the initiation of replication in new areas of the genome and block cell cycle progression to allow for DNA damage resolution ^1^. Insufficiencies in these S-phase responses and persistence of unreplicated DNA past S-phase, induce rescue mechanisms during mitosis, such as mitotic DNA synthesis (MiDAS) and anaphase-bridge resolution. Failure of all these pathways results in DNA breakage, chromosome mis-segregation and chromosomal rearrangements. Genome-wide, such unreplicated regions correlate with common fragile sites (CFS), which are chromosomal loci responsible for the majority of rearrangements found in human disease ^2^.

TRAIP (TRAF-interacting protein, also known as TRIP or RNF206) E3 ubiquitin ligase has recently been shown to play a role in a number of the above processes. TRAIP is essential for cell proliferation at an early stage of development in the mouse embryo ^3^ and CRISPR/Cas9-mediated deletion of TRAIP in a number of human cell lines is lethal ^4^. Moreover, homozygous mutations of the TRAIP ubiquitin ligase domain in humans lead to microcephalic primordial dwarfism ^5^.

At the cellular level, TRAIP has been shown to be essential for the appropriate repair of DNA damage caused by mitomycin C (MMC) and other inter-strand crosslinks (ICLs) generating drugs; camptothecin (CPT); UV and hydroxyurea (HU)^5–8^. TRAIP has also been reported to regulate mitotic progression: cells with downregulated TRAIP go through mitosis faster and with more chromosome segregation errors ^9–10^. TRAIP usually accumulates in nucleoli in cells and re-localizes to sites of DNA damage and replication stress ^5–7,11^. This co-localisation with PCNA upon DNA damage is mediated by a PCNA-interacting protein (PIP)-box motif located at the C-terminus of TRAIP ^6,7^. However, TRAIP was also shown to interact with unchallenged replication forks in *Xenopus egg* extract ^12^ and, in human cells, TRAIP has been shown to interact with nascent DNA during unperturbed S-phase through Nascent Chromatin Capture (NCC)and iPOND ^6,13,14^.

At the molecular level, TRAIP has been shown to orchestrate the response to ICLs in *Xenopus laevis egg* extract through the ubiquitylation of the eukaryotic replicative helicase (CMG complex, from <u>C</u>DC45/<u>M</u>CM2-7/<u>G</u>INS) as two replication forks converge at the ICL ^12^. In such a situation, short ubiquitin chains synthesized by TRAIP on CMGs promote the recruitment of NEIL3 glycosylase and unhooking of the ICL, whilst longer ubiquitin chains are required for CMG unloading by p97 segregase, allowing access for endonucleases and Fanconi anemia pathway proteins that perform ICL repair ^12,15,16^. Using the same model system, TRAIP has also been shown to act when the replisome encounters DNA-protein crosslinks (DPCs), which impair replication forks progression. In such a situation, however, TRAIP ubiquitylates not the CMG helicase within the blocked replisome, but the protein barrier itself ^17^. Moreover, in human cells, TRAIP was also shown to interact with RNF20-RNF40 ubiquitin ligase at double-strand breaks and affect ionizing-radiation induced monoubiquitylation of histone H2B ^18,19^, as well as the localization of BRCA1 interacting partner RAP80 to DNA double-strand breaks ^18^.

On the other hand, in mitosis, in *Xenopus leavis egg* extract, *C. elegans* embryos and mouse embryonic stem cells, we and others have shown that TRAIP ubiquitylates any replicative helicases left on chromatin from replication in S-phase, leading to their unloading by the p97 segregase ^20–23^. Such replisomes retained on chromatin until mitosis likely protect the DNA they are bound to, preventing access and subsequent DNA processing by nucleases. As a result, TRAIP-driven replisome disassembly in mitosis can lead to fork breakage and complex DNA rearrangements in *Xenopus egg* extract ^22^ and allows for MiDAS in human cells ^21^. Finally, TRAIP depletion was reported to lead to decreased stability of kinetochore-microtubule attachments and diminished spindle assembly checkpoint function through lowered MAD2 levels at centromeres ^9,24^.

With so many varied functions of TRAIP reported, the question arises as to which one of them is most crucial for cell viability, proliferation and prevention of microcephalic dwarfism? To answer these questions, we have generated auxin inducible degrons of TRAIP in the colon carcinoma HCT116 cell lines, which facilitate the degradation of TRAIP within 30 min of auxin (IAA) addition to the cell media. With these, we could confirm that upon rapid TRAIP degradation cells cease to proliferate, arrest in G2 stage of the cell cycle and undergo senescence. By further investigating the effect of TRAIP degradation in specific stages of the cell cycle we found that TRAIP plays its essential role in S-phase and that the lack of TRAIP results in the generation of DNA damage at sites of replication-transcription collisions.

## Materials and Methods

### Plasmid Construction

DONOR and CRISPR plasmids were constructed for the generation of conditional Auxin-Inducible Degron Cell lines as detailed in Natsume *et al.*, (2016) ^25^. Briefly, CRISPR-Cas9 was expressed using the pX330-U6-Chimeric_BB-CBh-hSpCas9 plasmid ^26^ (Addgene #42230) and targeted the C-terminus coding region containing the stop codon (5’-CTCACTGTTCTCACGACCAC-3’). Donor plasmids were generated following a published protocol^27^. Briefly, we cloned homology arms upstream and downstream of the CRISPR target region to pBluescript^25,27^ (about 500 bp each). After inverse PCR, a cassette containing mAID or mAID-Clover (mAC) with a selection marker was cloned to make a donor plasmid. Two donor plasmids were created: TRAIP-mAID and TRAIP-mAC, both containing Hygromycin resistance for selection.

### Generation of HCT116 TRAIP Conditional Degron Cell Lines

HCT116 cell lines constitutively expressing the *Oryza sativa* auxin-sensitive F-box protein (OsTIR1) ^25^ were transfected with the previously described DONOR and CRISPR plasmids. Parental cell lines were seeded to individual wells of a 6-well plate at a final working concentration of 1 × 10^5^ cells/ml. Transfection was conducted 2 days following plating using Fugene HD, OPTI-MEM, 200 ng/μl DONOR plasmid, and 200 ng/μl CRISPR plasmid. Control reactions were transfected with DONOR plasmids only. Approximately 24 hours later, cells were collected and diluted at different concentrations to 10 cm dishes. Selection was carried out 24 hours after dilution (100 μg/ml Hygromycin) and maintained continuously for 11-13 days. Once sufficient colonies were observed, 48 from each condition were isolated, grown in individual wells of a 96-well plate until confluent, and screened for bi-allelic gene insertion using genomic PCR. Adherent cells were washed twice with Dulbecco’s PBS (-) (D-PBS (-)) and treated with DirectPCR Lysis Reagent (Viagen: 0.5X DirectPCR Lysis Reagent, 20 mg/ml proteinase K) overnight at 55°C. Proteinase K activity was stopped through incubation of the 96-well plate at 85°C for 90 minutes in humid conditions. Genomic PCR was carried out using GoTaq HotStart Green Master Mix (Promega), as per manufacturer’s instruction (TRAIP F: AGATGTGGTGAGTGTGGCTTC; TRAIP R: GCTGCAGTGATCTCATTCTTTCT; mAID: ATCTTTAGGACAAGCACTCTTCTCC). Bi-allelic gene insertion was confirmed by DNA gel electrophoresis using 2% agarose gels. All cells were maintained in McCoys’s 5A medium supplemented with 10% Fetal Bovine Serum, 2 mM L-glutamine, 100 U/ml Penicillin and 100 μg/ml Streptomycin; grown at 37°C with 5% CO_2_.

### Detecting TRAIP Protein Levels

Immunoblotting was used to detect both TRAIP band shifts corresponding to the inserted tag size, and protein degradation. Whole cell extracts were generated by re-suspending harvested cell pellets in UTB extraction buffer (8 M Urea, 50 mM TRIS-HCl, 150 mM B-Mercaptoethanol) for 10 minutes on ice. The cell lysate was then sonicated (BioRuptor: 30 sec on, 30 sec off, 5 min cycles, medium power) and centrifuged at 15 000 RPM for 20 minutes to separate soluble and insoluble fractions. The soluble fraction was collected and mixed with 4X SDS-PAGE Loading Buffer (NuPAGE) to a final concentration of 1X. Approximately 50 μg of total protein content was run on 4-12% gradient SDS-PAGE gels (Invitrogen) and transferred to Nitrocellulose membranes for 90 minutes at 80 V. Transferred membranes were blocked (5% Milk in TBST) for 1 hour before being incubated in primary antibody at 4°C overnight. Membranes were washed 3X 10 minutes in TBST and incubated in secondary antibody for 2 hours at room temperature. The membranes were then washed 3X 10 minutes in TBST before being developed using ECL detection spray (Advansta WesternBright). Antibodies used for immunoblotting were as follows: Anti-TRAIP antibodies were kindly provided by Prof. N. Mailand ^6^ and used 1:300 in 5% Milk in TBST. Anti-β Actin loading controls used 1:5000 in 5% BSA in TBST (C4 anti-Actin HRP Santa Cruz).

### Drug Treatments

TRAIP degradation was induced through the addition of 500 μM Indole-3 acetic acid (IAA) to cell media, subsequently diluted 1:5 in the existing growth media to provide a final working concentration of 100 μM. In addition, the respective working concentrations of other inhibitor treatments used are described: 10 μM ATM inhibitor (Stratech KU-55933), 4 μM ATR inhibitor (Stratech AZD6738), 5 μM MK-1775 (Stratech), 100 nM Triptolide (Sigma-Aldrich T3652), 100 μM DRB (Sigma-Aldrich D1916), 0.4 μM Aphidicolin (A0781, Sigma)

### Cell Viability Assays

For colony forming assays, cells were diluted to a working concentration of 1000 cells/ml and plated to individual wells of a 6-well plate at different seeding concentrations (100, 250, 500, 750, 1000 cells/well). Auxin (IAA) was added 24 hours after plating, and cells incubated until sufficient colony formation was observed (approximately 12-14 days). Cell media was removed, and colonies stained using Methylene Blue staining (2% methylene blue in 50% ethanol) buffer for 5 minutes at room temperature. Colonies were rinsed with H_2_O and dried overnight. For quantification, the percentage colony forming efficiency was calculated by dividing the number of counted colonies by the original number of cells plated, allowing comparison between different seeding concentrations. To then allow comparison between different cell lines, the calculated percentage colony forming efficiency was normalised to a control cell line (HCT116 CMV-OsTIR1 only, - IAA; Efficiency in test clone / Efficiency in Control). For cell proliferation assays, cells were diluted to 1 × 10^4^ cells/ml and plated into 60-mm dishes (4 × 10^4^ starting concentration). Auxin (IAA) was added 24 hours later, and cells grown for a further 72 hours. At each timepoint (24, 48, 72 hrs post auxin (IAA) treatment) cells were harvested and the total cell numbers counted using a COUNTESS cell counter. Cell counts were normalised to the seeding concentration. In addition, following cell counting, the remainder of the samples were prepared for Flow Cytometry as detailed.

### Cell Death Assays

Cell Senescence was detected using the Cell Meter Senescence Activity Assay (AAT Bioquest). Cells were plated for proliferation curves as described. At each timepoint, cells were harvested and washed with PBS. Cell pellets were resuspended in Xite Green β-D-galactopyranoside solution for 45 minutes at 37°C. Stained cells were washed again in PBS, resuspended in 500 μl Assay Buffer, and analysed using a Flow Cytometer. Senescent cells were detected using the FITC 488 channel, with unstained controls utilised to distinguish between positive and negative cell populations. Apoptotic cells were detected using the Annexin V Apoptosis Detection Kit (Invitrogen). Cells were grown and harvested as detailed. The resulting pellet was resuspended in 1X binding buffer supplemented with AF488 fluorochrome coated Annexin V. Samples were incubated for 15 minutes at room temperature and washed in Binding Buffer. Stained cells were then resuspended in 200 μl Binding buffer supplemented with 50 μg/ml Propidium Iodide and 50 μg/ml RNase A. Cells analysed using a flow cytometer, with apoptotic cells detected using the FITC 488 channel.

### Cell Synchronisation

For G1 cell synchronisation, asynchronous cells were treated with 20 μM Lovastatin (Acros Organics) for 24 hours. To then release cells from the G1 arrest, cells were washed 3X in prewarmed growth media before fresh media supplemented with 2 mM Mevalonic Acid (Sigma-Aldrich) was added. Cells entered S-phase approximately 12 hours post release. To arrest cells in G2 stage of the cell cycle, asynchronous cells were treated with 9 μM RO-3306 (Merck Life Sciences) for 16 hours. G2 release was achieved through washing the cells 3X in pre-warmed media before fresh growth media was added. Released cells enter mitosis approximately 30-60 minutes post release. To sequentially arrest cells in G1 and G2, cells were treated as described for G1 cell cycle arrest. Approximately 8 hours after G1 release, 9 μM RO-3306 was added and the cells incubated for 16 hours to facilitate G2 arrest. Cells were released as detailed.

### Immunofluorescence

Asynchronous cells were seeded onto pre-sterilised 20 mm glass cover slides placed into each well of a 6-well plate. Any respective drug treatments were carried out as described. To label S-phase cells, 10 μM EdU treatments were included 60 minutes prior to cell fixation. To fix cells, the growth media was removed, and the cells washed once in D-PBS (-). Washed cells were fixed using 4% Paraformaldehyde in PBS for 15 minutes at room temperature. Cells were then permeabilised in 0.5% TritonX-100 in PBS for 5 minutes at room temperature and washed twice. If required, EdU Click-IT was carried out as per the manufacturer’s protocol (Invitrogen) to detect S-phase cells prior to antibody staining. Antibody staining was subsequently carried out: 100 μl of primary antibody (in Washing Buffer: 5% BSA 0.1% Tween-20 in PBS) was added dropwise to cover the glass cover slip and incubated for 2 hours at room temperature. Cells were then washed 3X in washing buffer and 2X in D-PBS (-). Secondary antibody solution was added analogous to primary antibody and incubated for 2 hours at room temperature in the dark. Antibodies used are described: Mouse anti-Ser139 γ-H2AX (Sigma-Aldrich JBW301; 1:1000), Rabbit anti-53BP1 (Novus Biologicals NB100-904; 1:1000), Mouse anti-Mitosin (BD Biosciences 610768; 1:300), Rabbit anti-P-Histone H3S10 (Cell Signalling 9701; 1:1000), AF488 anti-mouse secondary (Invitrogen A32723; 1:1000), AF555 anti-rabbit secondary (Invitrogen A21428; 1:1000). Stained cells were washed as detailed before being mounted onto glass slides using DAPI mounting media (Fluoroshield). Slides were dried at room temperature before being imaged using a Leica DM600 Widefield Fluorescent microscope. All images were exported as raw grayscale TIFs, analysed using CellProfiler (v4.0.6) and ImageJ (v2.1.0).

### Flow Cytometry

Three types of flow cytometry experiment were carried out: un-extracted cells, BrdU detection, and extracted cells. For un-extracted cells, following the required experimental procedures (e.g., proliferation curves) cells were harvested and fixed in 70% ethanol in PBS for 16 hours at - 20°C. Following fixation, cells were washed twice in washing buffer (5% BSA, 0.1% Tween-20, PBS) and antibody staining carried out if required. Briefly, washed cells were resuspended in 100 μl primary antibody in washing buffer and incubated at room temperature for 1 hour, rocking to prevent cells from settling. The cells were washed twice in washing buffer and resuspended in 100 μl secondary antibody in washing buffer for 1 hour at room temperature in the dark. Stained cells were washed 1X in washing buffer and 2X in D-PBS (-) before being resuspended in either Hoescht Staining Buffer (5 μg/ml Hoescht 33582, PBS) or Propidium Iodide Staining Buffer (50 μg/ml Propidium Iodide, 50 μg/ml RNase A, PBS). For BrdU detection, 10 μM of BrdU was added to the growth media 1 hour prior to harvesting. Cells were collected and fixed in ethanol as described. Fixed cells were washed once in PBS before being resuspended in 1 ml 2 M HCL supplemented with 0.1 mg/ml Pepsin for 20 minutes. Cells were then washed, and antibody staining carried out as described. To explore the replisome binding pattern on chromatin, cells were extracted using CSK buffer (25 mM HEPES pH 7.4, 50 mM NaCl, 3 mM MgCl_2_, 300 mM Sucrose, 0.5% Triton X-100, 1X complete protease inhibitors) to remove soluble fractions. The protocol used to extract cells has been described elsewhere (Forment & Jackson, 2015). Antibodies used are as follows: anti-Histone H3 S10 phosphorylation (Cell Signalling 9701; 1:500), anti-MCM7 (Santa Cruz 9966; 1:500), anti-BrdU (Sigma-Aldrich, clone B44; 1:5), AF488 anti-mouse secondary (Invitrogen A32723; 1:1000), AF555 anti-rabbit secondary (Invitrogen A21428; 1:1000), AttoN647 anti-rabbit secondary (Sigma-Aldrich 40839; 1:500). Cells were analysed using a Beckman Cytoflex instrument. Data analysis was carried out using FlowJo (v10.7.1).

### ChIP-Sequencing

For ChIP-sequencing, HCT116 TRAIP degron cells were diluted to 2 × 10^5^ cells/ml and seeded onto 20 cm dishes (2X per condition). Plated cells were arrested in G1 as described, treated with auxin (IAA) to degrade TRAIP, and released into S-phase. 16 hours after release, cells were harvested and the resulting cell suspension supplemented with 1% Formaldehyde (Sigma-Aldrich) for 10 minutes at room temperature. Formaldehyde crosslinking was quenched using 125 mM Glycine (Sigma-Aldrich). Fixed cells were pelleted (400 g, 3 minutes, 4°C) and the pellet washed 2X with ice-cold PBS. Cell extraction was then carried out through sequential incubations in ChIP lysis buffer (5 mM HEPES pH 8.0, 85 mM KCL, 0.5% NP40) and ChIP nuclear lysis buffer (50 mM Tris-HCL pH 8.0, 10 mM EDTA pH 8.0, 1% SDS) for 15 and 30 minutes on ice, respectively. Cell lysates were divided into equal aliquots and sonicated (30 amplitude, 15 sec on, 25 sec off, 12 cycles). Sufficient sonication (resulting in 300-500 bp DNA fragments) was confirmed through DNA gel electrophoresis. Chromatin immunoprecipitation was carried out using Protein A Dynabeads conjugated to approximately 1 μg of S139 rabbit γ-H2AX (Abcam 29893) or Rabbit H2AX (Merck Millipore 07627) antibodies, as detailed in Wang *et al*., (2021) ^28^. Immunoprecipitation was validated by quantitative PCR using primers targeting the actin housekeeping gene (Forward: CATGTACGTTGCTATCCAGGC, Reverse: CTCCTTAATGTCACGCACGAT).

Library preparation was carried out using NEBNext Ultra II DNA Library Preparation Kit for Illumina NEB, as per manufacturer’s instruction. The prepared libraries were sequenced using single-end sequencing with a High-75 kit, using Illumina NexSeq instruments. γ-H2AX ChIP-Seq, precision run-on sequencing (PRO-Seq) ^29^ and RNA Pol II chromatin immuno-precipitation sequencing ^30^ in HCT116 cells were aligned to the hg38 genome using Bowtie 2 v.2.4.2 on the online platform Galaxy (https://usegalaxy.org; ^31^). γ-H2AX ChIP-Seq peaks were called in the +IAA treated sample against the γ-H2AX ChIP-Seq in its -IAA control using MACS2 v.2.1.1 with parameters as detailed in ^28^. The bedtools intersect intervals function on the online platform Galaxy was used to identify the conserved γ-H2AX ChIP-Seq peaks in the two repeats. The distance between the γ-H2AX ChIP-Seq peak or the TSS with lowest γ-H2AX fold change +IAA/-IAA was calculated using the position of the origins of replication in HCT116 cells provided by Dr Daigaku ^32^. The PRO-Seq dataset was used to determine the reciprocal direction between the oncoming replication fork and gene transcription (head-to-head or codirectional), as well as whether the TSS was the first, second or third and above transcribed gene encountered. Replication timing for the γ-H2AX ChIP-Seq peak was derived analysing replication timing in HCT116 cells from ^33^. The read coverage profiles were generated using the computational environment EaSeq, normalizing the γ-H2AX ChIP-Seq file to the H2AX ChIP-Seq file with the function “average” ^34^.

To identify the list of transcribed genes in HCT116 cells, the counts for each gene were computed by featurecounts on the online platform Galaxy using the annotation of the GENCODE genes (GRCh38.p10) on an ENCODE polyA RNA-Seq hg38 aligned file (ENCFF823JEV). Read per kilobase per million (RPKM) were calculated over each gene and genes that had an RPKM > 1 were considered as transcribed.

The levels of antisense to sense transcription were calculated using the PRO-Seq and RNA Pol II ChIP-Seq datasets using the function “quantify” in EaSeq, for the antisense −1000 bp to the TSS, and for the sense from the TSS to +1000bp. For the PRO-Seq, as this was strand-specific antisense transcription levels were specifically measured on the strand opposite to the sense transcription. Heatmaps were generated with the function “HeatMap” of EaSeq around TSS +/- 2500 bp. To identify the genes with the lowest increase in γ-H2AX ChIP-Seq levels following TRAIP depletion, the normalised levels of γ-H2AX ChIP-Seq signal to H2AX was calculated across the TSS +/- 1000 bp of all transcribed genes in both the repeats and averaged for each gene. Then gene TSS were sorted by the fold change in γ-H2AX ChIP-Seq levels in the +IAA compared to the -IAA.

### Proximity-Ligation Assays

Proximity Ligation Assays were carried out using the DuoLink PLA Kit. The provided manufacturer’s instruction was optimised for use on cell suspensions. Parental cell lines were diluted to 2 × 10^5^ cells/ml and seeded to 60-mm tissue culture dishes. Cells were arrested in G1, where auxin (IAA) was added, and released into S-phase. Approximately 12 hours following cell cycle release, 10 μM EdU was added for 20 minutes to label nascent DNA. Cells were then harvested and extracted using CSK buffer as described previously. Extracted, permeabilised cells were subjected to the Click-It reaction to conjugate biotin to incorporated EdU (Invitrogen, as per manufacturer’s protocol). Cells were washed 2X in PLA washing buffer and resuspended in 50 μl of each primary antibody made up in washing buffer (elongating RNA Polymerase II: Mouse anti-Serine 5 CTD, Cell Signalling; Rabbit anti-biotin, Bethyl Laboratories A150-109A) overnight at 4°C. Following primary antibody incubation, cells were washed 3X in PLA wash buffer A and incubated in 30 μl secondary antibody solution (6 μl ‘+’, 6 μl ‘-’ 18 μl 3% FBS in PBS) for 100 minutes at 37°C. Cells were washed again 3X in washing buffer A and resuspended in 30 μl Ligation Buffer, incubated for 60 minutes at 37°C. Any interacting PLA probes were then amplified using a rolling circle assay. Washed cell pellets were resuspended in PLA Amplification buffer for 100 minutes at 37°C. Finally, cells were washed 2X in washing buffer B, resuspended in 0.01X washing buffer B diluted in distilled H_2_O. Diluted cell suspensions were added to 20 mm cover slips and centrifuged to adhere cells to cover slips (400 g, 5 min). Slides were mounted using DuoLink PLA DAPI mounting media. Cells were analysed using a Leica DM600 widefield microscope.

#### *XENOPUS LAEVIS* METHODS

##### Inhibitors and Recombinant Proteins

EcoR1 (R6011, Promega) was purchased at stock 12 U/μl and added to the extract at 0.05 U/μl, Aphidicolin (A0781, Sigma) was dissolved in DMSO at 8 mM and added to the extract along with demembranated sperm nuclei at 40 μM. Caffeine (C8960, Sigma) was dissolved in water at 100 mM and added to the extract along with demembranated sperm nuclei at 5 mM. MLN4924 (A01139, Active Biochem) was dissolved in DMSO at 20 mM and added to the extract 15 minutes after addition of sperm nuclei at 10 μM.

Recombinant His-tagged *Xenopus laevis* CyclinA1 NΔ56 (pET23a-X.l cyclin A1 NΔ56), was expressed and purified as previously described ^20^. CyclinA1 NΔ56 was used at a final concentration of 826 nM in the egg extract to drive the extract into mitosis.

##### Antibodies

anti-PCNA (P8825) was purchased from Sigma; anti-TRAIP (NBP1-87125) was purchased from Novus Biologicals; anti-P-Chk1 (S345) from Cell Signalling, γ-H2AX (4418-APC-020, Trevigen). Affinity purified anti-Cdc45, anti-Psf2 ^35^, anti-Mcm7 ^20^ and anti-GINS antibody ^36^ were previously described.

Recombinant *Xenopus laevis* SUMO-TRAIP was purified as previously described ^20^ and was also used for raising antibodies in rabbits. The resulting antibody sera was purified in-house against the purified antigen. The specificity of this new antibody is presented in Supplementary Figure 10A.

##### *Xenopus laevis* Egg Extract Preparation

Metaphase II arrested egg extracts were prepared as previously described ^37^.

##### DNA Synthesis Assay

Interphase *Xenopus laevis egg* extract was supplemented with 10 ng/μl of demembranated *Xenopus* sperm nuclei and incubated at 23°C for indicated times. Synthesis of nascent DNA was then measured by quantification of radiolabelled α^32^P-dATP (NEG512H250UC, Perkin Elmer) incorporation into newly synthesised DNA, as described before ^37^.

##### Chromatin Isolation Time-course

Interphase *Xenopus laevis* egg extract was supplemented with 10 ng/μl of demembranated sperm DNA and subjected to indicated treatments. The reaction was incubated at 23°C for indicated length of time when chromatin was isolated in ANIB100 buffer (50 mM HEPES pH 7.6, 100 mM KOAc, 10 mM MgOAc, 2.5 mM Mg-ATP, 0.5 mM spermidine, 0.3 mM spermine, 1 μg/ml of each aprotinin, leupeptin and pepstatin, 25 mM β-glycerophosphate, 0.2 μM microcystin-LR and 10 mM 2-chloroacetamide (Merck) as described previously ^37^. To study mitosis, the interphase extract was supplemented with MLN4924 and allowed replication to complete. Once completed extract was treated with CyclinA1NΔ56.

During the chromatin isolation procedure, a sample without addition of sperm DNA (no DNA) is processed in an analogous way, usually at the end of the time course, to serve as a chromatin specificity control. The bottom of the PAGE gel on which the chromatin samples were resolved was cut off and stained with Colloidal Coomassie (SimplyBlue, Life Technologies) to stain histones which provide loading controls and indications of sample contamination with egg extract (cytoplasm).

##### Nuclei Isolation for Chk1 Phosphorylation

The nuclei isolation was performed as previously described ^38^.

##### Immunodepletion

TRAIP immunodepletions were performed using Dynabeads Protein A (10002D, Life Technologies) coupled to *Xenopus* TRAIP antibodies raised in rabbit and affinity purified or nonspecific rabbit IgG (I5006, Sigma). The TRAIP antibodies were coupled at 600 μg per 1 ml of beads. Effective immunodepletion required 2 rounds of 1 h incubation of egg extract with antibody coupled beads at 50% beads ratio (e.g. 2 rounds of 100 μl of egg extract incubated with 50 μl of coupled beads). The immunodepletion was performed as described before ^38^.

##### Statistical Analysis

All statistical analysis except from ChIP-Sequencing data was carried out using RStudio (v 1.0.153). The imported data was first subject to normality testing through qqplots to determine the appropriate statistical testing method. All plots were created using ggplot2 and the plug-in ggpubr (^39^). Two biological repeats for each γ-H2AX and H2AX ChIP-Seq in -IAA and +IAA were analysed, with repeats assessed for correlation before being combined. Student t test and Mann-Whitney t test were calculated using the software Prism (GraphPad).

## Results

To investigate the role of TRAIP during the unperturbed eukaryotic cell cycle, we established conditional auxin-inducible degron (AID) cell lines ^25,27^. As modifying the N-terminus of TRAIP was reported to impact on TRAIP localization in the cell^7^, endogenous TRAIP was tagged C-terminally in a HCT116 cell line expressing OsTIR1, an E3 ligase component recognizing the degron, with either a mini-auxin inducible degron (mAID) tag, or an mCLOVER-fused variant (mAC); henceforth referred to as TRAIP-mAID and TRAIP-mAC, respectively (Supplementary Figure 1A and 1B). Bi-allelic tagging was verified by PCR amplification from the genomic TRAIP locus (Supplementary Figure 1C). It was also confirmed on protein level by disappearance of endogenous untagged TRAIP protein from the whole cell extracts prepared using each degron cell line (Supplementary Figure 1D). Bi-allelic modification of TRAIP with degron tags did not affect cell proliferation and fitness (Supplementary Figure 1E). Activation of the AID system, through the addition of Indole-3-acetic acid (auxin, IAA) to the cell growth media ensured rapid protein degradation in approximately 30 minutes (Figure 1A, Supplementary Figure 1F).

**Figure 1.**
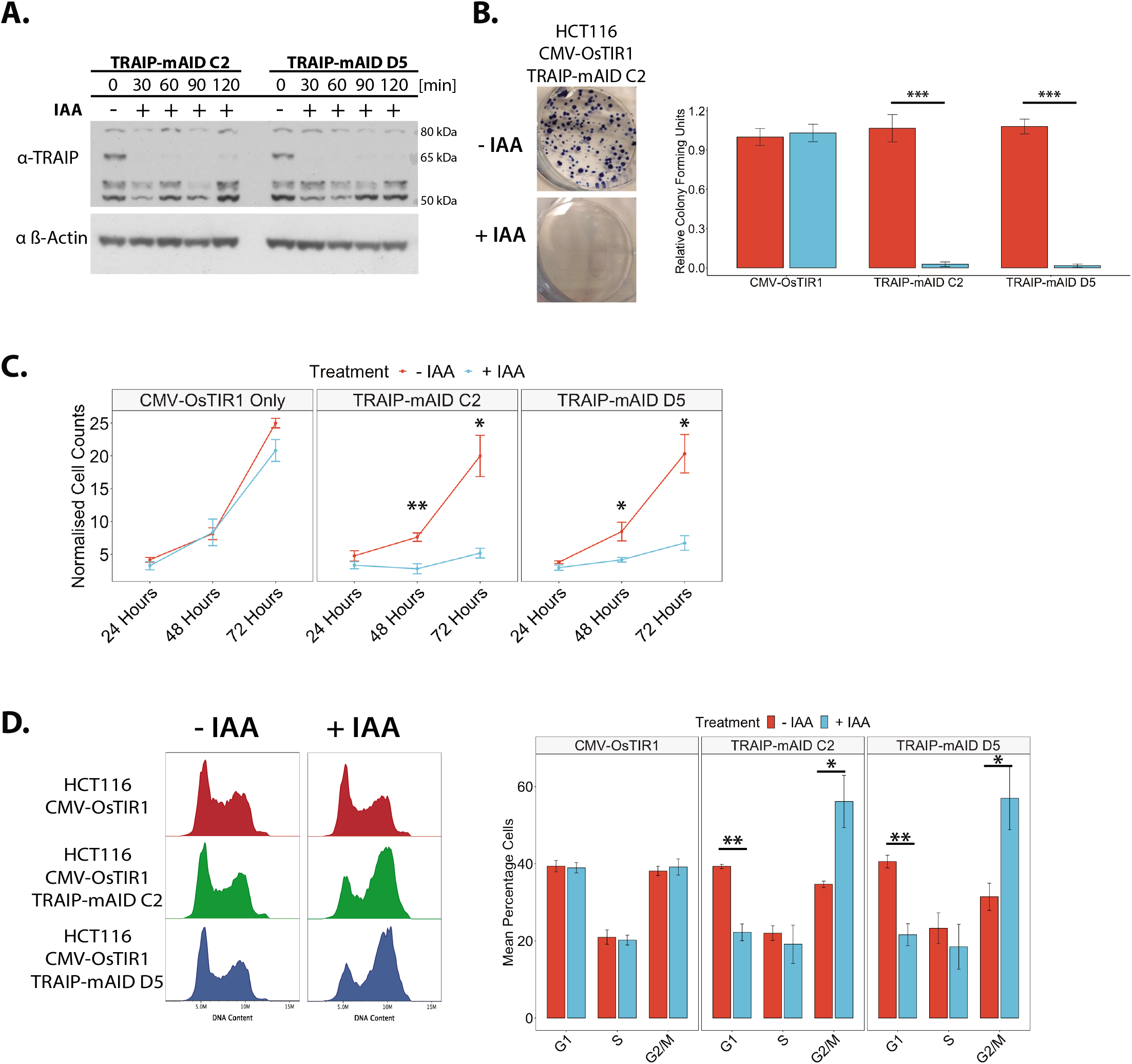
Rapid auxin induced degradation of TRAIP-mAID leads to G2 cell cycle arrest and inhibition of cell proliferation. **(A)** TRAIP-mAID is degraded within 30 min of auxin (IAA) addition. An asynchronous cell culture of HCT116 TRAIP-mAID cells was supplemented with auxin (IAA) and samples taken at indicated times. The level of TRAIP-mAID was assessed by western blotting with TRAIP antibody. Two independent clones of TRAIP-mAID are presented. **(B)** TRAIP degradation leads to the loss of cell viability. Cells were grown in presence or absence of auxin (IAA) and resultant colonies visualised. Example colony assay (left) and quantification of 3 biological repeats (right). Pairwise hypothesis testing was conducted using Mann-Whitney U Tests, with significance values indicated on the plot (TRAIP-mAID C2: p < 0.001; TRAIP-mAID D5: p < 0.001) **(C)** TRAIP degradation ceases cell proliferation within first cell cycle without TRAIP. TRAIP-mAID cells were grown in optional presence of auxin (IAA) and cell proliferation counted over the 72 h. Quantification of n=3 for two independent TRAIP-mAID clones. Data is shown as the mean +/- SEM. Pairwise statistical comparison were conducted at each timepoint using t.tests, with significance indicated on the plot (TRAIP-mAID C2: 48 h – p = 0.00940, 72 h – p = 0.0363; TRAIP-mAID D5: 48 h – p = 0.04, 72 h – p = 0.0305). **(D)** TRAIP degradation leads to cell cycle arrest with G2/M DNA content. TRAIP-mAID cells (two independent clones) were grown for 24 h in optional presence of auxin (IAA) and analysed for their DNA content by FACS. Example FACS plots (left) and quantification of number of cells in different stages of the cell cycle (mean +/- SEM) over 3 independent experiments for 2 independent clones (right) are shown. Pairwise comparisons were carried out using t.tests, significance is indicated on the plot (TRAIP-mAID C2: G1 – p = 0.0120, G2/M – p = 0.04; TRAIP-mAID D5: G1 – p = 0.00874, G2/M – p = 0.04).

### TRAIP activity is essential for cell proliferation

We first set out to understand the global consequences of TRAIP depletion for cell viability, proliferation and cell cycle progression. Colony assays were used to assess the respective impact of auxin (IAA) treatment itself, and TRAIP degradation, on cell viability. Long-term auxin (IAA) treatment (7-14 days) led to abrogation of colony growth in all the conditional TRAIP degron cell lines (Figure 1B; Supplementary Figure 2A). Importantly, no reductions in the overall colony forming ability were found when auxin (IAA) was added to control cell lines, indicating that the drug treatment itself was not cytotoxic. These data confirmed that TRAIP was an essential protein as previously described ^3,4^. To investigate the underlying reason for the observed growth inhibition, we next investigated the effects of shorter periods of protein degradation (24-72 Hrs) on cell proliferation and cell cycle progression. Cells lacking TRAIP exhibited clear defects to cell growth at all timepoints tested (Figure 1C, Supplementary Figure 2B). Furthermore, flow cytometric analysis of representative samples taken at each timepoint revealed the reduction in the number of cells in G1 stage of the cell cycle, as well as the accumulation of TRAIP-depleted cells with G2/M DNA content (Figure 1D, Supplementary Figure 2C). Altogether, these results show that our TRAIP degron cell lines exhibit phenotypes analogous to those observed previously upon downregulation of TRAIP by siRNA or in patients’ cells with impaired ubiquitin ligase activity of TRAIP ^5–7^ and thus provide a useful tool for the determination of the cell-cycle stage specific, essential function of TRAIP.

### TRAIP depleted cells accumulate in G2 stage of the cell cycle

As the underlying mechanism of cell death in cells depleted of TRAIP has not been previously determined, we decided first to investigate the mechanism by which TRAIP depleted cells stop proliferating. We first determined in which cell cycle stage cells lacking TRAIP were accumulating: G2 or mitosis. Quantification of the total proportion of cells displaying phosphorylation of Histone H3 on S10 (marker of mitosis) by flow cytometry showed no differences to the numbers of mitotic cells following TRAIP degradation (24 h auxin (IAA) treatment) (Supplementary Figure 3A). The same was confirmed by mitotic indexing experiments (Supplementary Figure 3B). To specifically visualise G2 population of cells, cells were treated with auxin (IAA) for 24 hours to degrade endogenous TRAIP and stained for three markers of G2/M progression: DAPI for condensing chromosomes, Mitosin (CENPF) for G2 and mitosis, and the S10 phosphorylation of Histone H3 for mitosis (Figure 2A). A significant increase was found in the proportions of cells found to be in G2 stage of the cell cycle following treatment with auxin (IAA) and TRAIP degradation (Figure 2A, no WEE1i), while no differences again were found when considering those cells in either early or late mitosis (Supplementary Figure 3C). Finally, the G2 cell cycle accumulation was rescued through inhibition of the G2 checkpoint kinase WEE1 using MK-1775 inhibitor (Figure 2A and Supplementary Figure 3C). We conclude therefore, that the loss of TRAIP results in a G2 cell cycle arrest mediated by WEE1.

**Figure 2.**
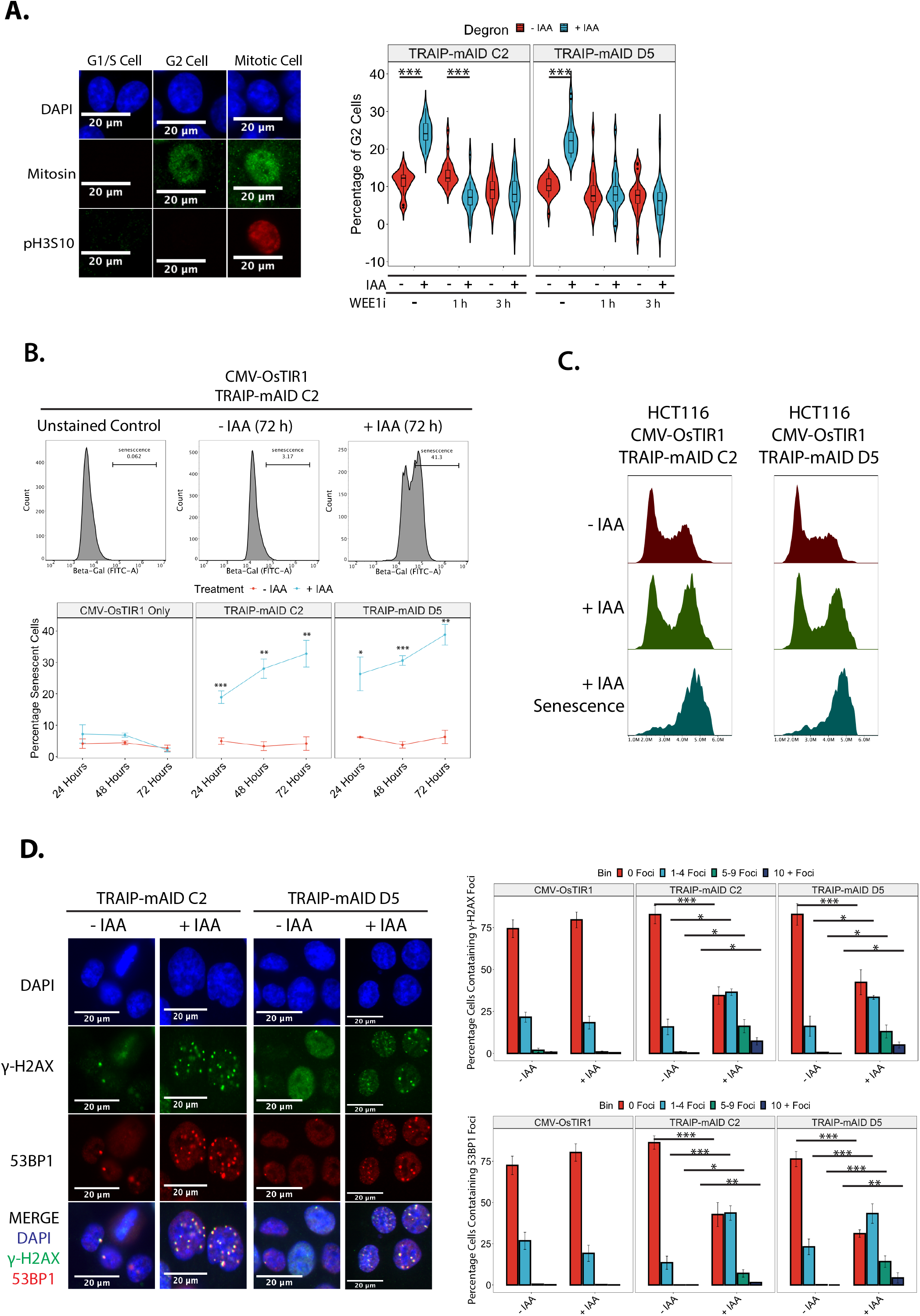
TRAIP degradation leads to DNA damage, G2 cell cycle arrest and cell senescence. **(A)** TRAIP degradation leads to accumulation of cells in G2 stage of the cell cycle. TRAIP-mAID cells (two independent clones) were grown for 24 h in optional presence of auxin (IAA), and optional addition of WEE1i for indicated times, and analysed for accumulation of cells in G2 (Mitosin staining) or in mitosis (pH3S10 staining). Quantification of 3 independent experiments. Data depicted as violin plots, displaying boxplots highlighting the median and corresponding interquartile ranges. Statistical analysis was carried out using one-way ANOVA and subsequent post-hoc testing, significance is indicated on the plots (ANOVA: p < 0.0001. Tukey’s post-hoc tests: all indicated comparisons on the plot p < 0.0001). **(B)** TRAIP-mAID cells were grown for 24-72 h in optional presence of auxin (IAA) and the level of senescent cells analysed by FACS measurement of senescence marker β-galactosidase. Example FACS plots (top) and quantification of 3 independent experiments in 2 independent clones (bottom). Quantified data is shown as the mean +/- SEM. Pairwise hypothesis testing was carried out using t.tests, with significance indicated on the plot (TRAIP-mAID C2: 24 h - p = 0.00118, 48 h - p = 0.00651, 72 h p = 0.00926; TRAIP-mAID D5: 24 h - p = 0.03, 48 h - p = 0.00031, 72 h - p = 0.0022). **(C)** TRAIP depleted cells go into senescence while in G2 stage of the cell cycle. TRAIP-mAID cells were grown for 72 h in optional presence of auxin (IAA). β-galactosidase staining was combined with DNA content analysis by FACS. β-galactosidase positive cells have G2/M DNA content. **(D)** Increased level of DNA damage is observed upon TRAIP degradation. TRAIP-mAID cells were grown in optional presence of auxin (IAA) for 24 h and stained with markers of DNA damage response (γH2AX and 53BP1). Example of immunofluorescent visualisation of DNA damage (left) and quantification of percentage of cells containing increasing numbers of γH2AX and 53BP1 foci over 3 independent experiments (right). Significance testing carried out using oneway ANOVA and subsequent Tukey’s post-hoc testing (TRAIP-mAID C2: ANOVA – p < 0.001; Tukey’s post hoc test – 0 Foci: p = < 0.001; 1-4 Foci: p < 0.001; 5-9 Foci: p = 0.0168; 10 + Foci: p = 0.0059. TRAIP-mAID D5: ANOVA - p < 0.001; Tukey’s post hoc test - 0 Foci: p < 0.001; 1-4 Foci: p < 0.001; 5-9 Foci: p = 0.0046.

### G2-arrested TRAIP-depleted cells exit the cell cycle through senescence

Analyses of each degron cell line treated for 24 h with auxin (IAA) also revealed an increase in the size of nuclei and overall cell size (Supplementary Figure 4A and B). Intriguingly, increased cell size, reductions to cell proliferation and cell cycle accumulation are all hallmarks of cell cycle exit via senescence^40^. To explore this further, we used flow cytometry to detect the senescence marker β-galactosidase. Indeed, following as little as 24 h of auxin (IAA) treatment, cells were positive for this marker of cell senescence (Figure 2B). Subsequent comparison of the positive β-galactosidase cell population with total DNA content revealed the specific activation of senescence in the accumulating G2/M cells (Figure 2C). In contrast, we have detected no increase in the proportion of apoptotic cells (Supplementary Figure 4C and D) and we observed no increase in cells exhibiting mitotic catastrophe phenotypes (data not shown).

### G2 arrest in TRAIP-depleted cells is a result of DNA damage checkpoint activation

G2 cell cycle arrest is often a consequence of DNA damage checkpoint activation ^41^ and TRAIP was previously linked to numerous DNA damage repair pathways. To check whether DNA damage accumulation upon degradation of TRAIP may be indeed responsible for the observed G2 arrest, cells were treated with auxin (IAA) for 24 hours and stained for the general DNA damage marker γ-H2AX (S139 phosphorylation on H2AX) as well as the double-stranded DNA break-specific marker 53BP1. This revealed a general, but subtle, accumulation of DNA damage foci in the two independent TRAIP-mAID cell lines (Figure 2D), indicating that TRAIP is important for the maintenance of genomic stability. Moreover, inhibition of ATR, but not ATM, was able to rescue the observed G2 cell cycle arrest (Supplementary Figure 5A) suggesting that the damage is created in S-phase during DNA replication. We conclude therefore, that the loss of TRAIP results in creation of DNA damage, checkpoint activation and the accumulation of cells in G2 stage of the cell cycle. As a result of this, cells enter senescence and stop proliferating.

### TRAIP is essential for S-phase progression

To understand specifically when during the cell cycle TRAIP was essential, we combined the rapid nature of the AID system with established cell synchronisation techniques. Due to ATR-dependent checkpoint activation in TRAIP depleted cells and the known roles of TRAIP in S-phase, we first investigated whether TRAIP was essential during S-phase. Cells were arrested in G1 and auxin (IAA) was added for the last hour of G1 arrest. Following release from G1 arrest, timepoints were taken to assess the relative impact of TRAIP degradation on the cell cycle using the previously described G2 cell cycle arrest as a readout. When cells progressed through S-phase and into G2 lacking TRAIP, G2 cell cycle accumulation could be observed (Figure 3A). To verify whether TRAIP is required specifically during the G2 stage of the cell cycle, TRAIP was optionally degraded 15 hours post release from G1 arrest, when cells were in late S-phase. When auxin (IAA) was added during late S-phase, no G2 cell cycle accumulation could be detected (Figure 3B). These data indicated that TRAIP was essential specifically for mechanisms existing during S-phase.

**Figure 3.**
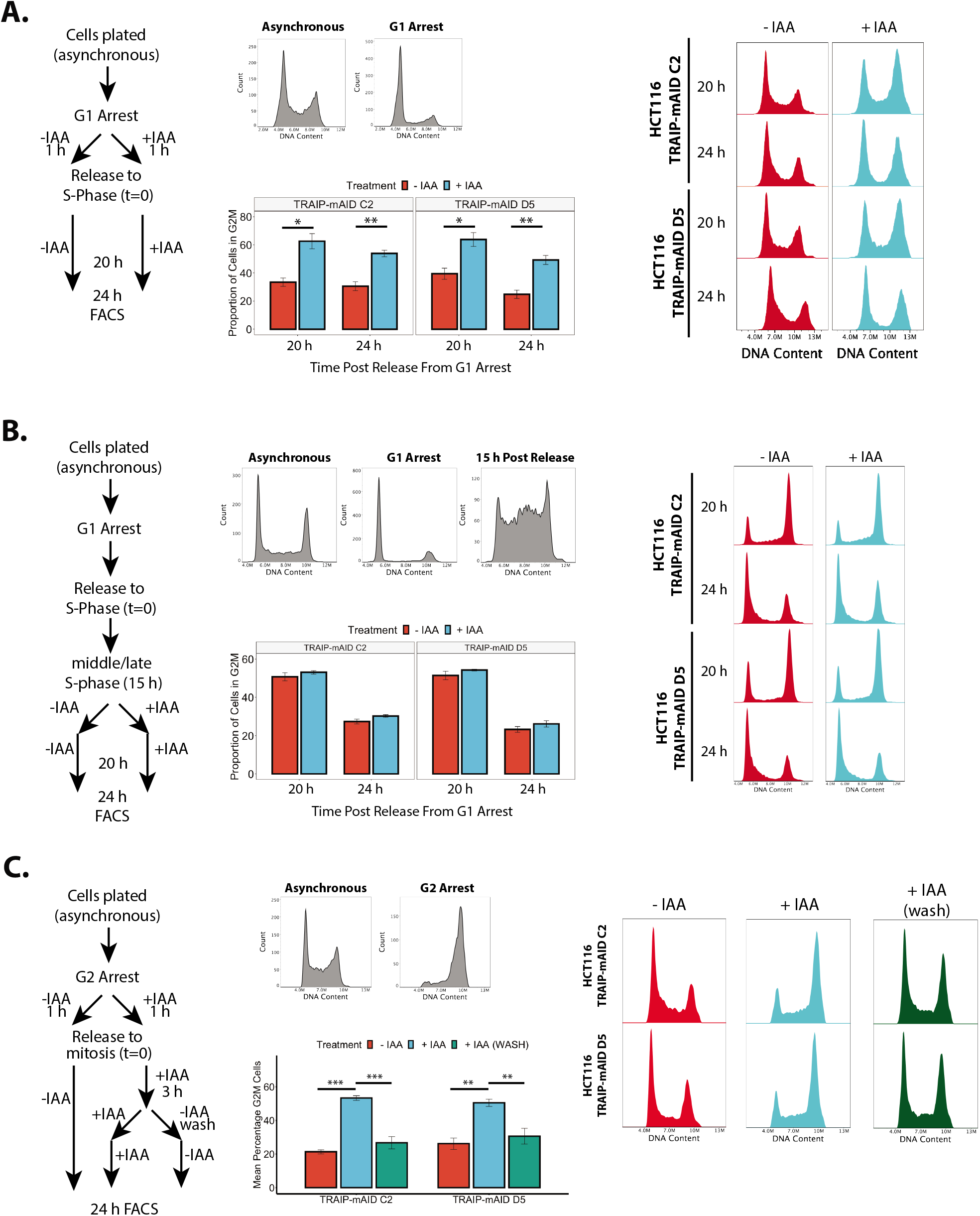
TRAIP is essential for S-phase progression during unperturbed cell cycle. **(A)** Degradation of TRAIP specifically just before S-phase leads to G2 cell cycle arrest. TRAIP-mAID cells were arrested in G1 stage of the cell cycle and TRAIP degraded before release of cells into S-phase. Cell cycle progression was analysed by FACS at 20 and 24 h post release. The experimental set up (left), example of cell cycle profiles at the time of synchronisation (middle, top), examples of FACS profiles at 20 and 24 h post release (right) and quantification of cells with G2/M DNA content (mean +/- SEM) over 3 independent experiments with 2 independent clones (middle, bottom). Pairwise hypothesis testing conducted using t.tests, with significance indicated (TRAIP-mAID C2: 20 h G1 – p = 0.0288, 24 h G1 – p = 0.0171, 20 h G2/M – p = 0.00514, 24 h G2/M – p = 0.00514; TRAIP-mAID D5: 20 h G1 – p = 0.00825, 24 h G1 – p = 0.0196, 20 h G2/M – p = 0.0198, 24 h G2/M – p = 0.00517. **(B)** Lack of TRAIP in G2 stage of the cell cycle does not lead to G2 cell cycle arrest. TRAIP-mAID cells were arrested in G1 stage of the cell cycle, released into S-phase and auxin (IAA) added in middle/late S-phase. Cell cycle progression was analysed by FACS at 20 and 24 h post cell release into S-phase. All analyses presented as in (A). **(C)** Mitotic functions of TRAIP do not contribute towards problems leading to G2 cell cycle arrest. TRAIP-mAID cells were arrested in G2 stage of the cell cycle and TRAIP degraded before release of cells into mitosis. After release auxin (IAA) was optionally washed off 3 h after release, so that cells progressed through next S-phase with or without TRAIP. Cell cycle progression was analysed by FACS at 24 h post release from G2 arrest. All analyses presented as in (A). Any difference between treatment groups was determined using one-way ANOVA and post-hoc testing, significance is summarised on the plot (one-way ANOVA: p < 0.0001; Tukey’s post-hoc testing, TRAIP-mAID C2: - IAA vs + IAA, p < 0.001; - IAA vs + IAA WASH, p = 0.800; + IAA vs + IAA WASH, p < 0.001. TRAIP-mAID D5: - IAA vs + IAA, p = 0.00107; - IAA vs + IAA WASH, p = 0.898; + IAA vs + IAA WASH, p = 0.00554).

Finally, TRAIP was previously reported to have many functions also during mitosis ^9,10,20–23^. To determine whether we could observe that TRAIP regulates mitotic progression in our cells, we arrested cells specifically in G2 stage of the cell cycle with RO-3306, degraded TRAIP, released cells to mitosis and followed chromosomal condensation and segregation (Supplementary Figure 5B). We could observe that irrespectively of TRAIP degradation, cells condensed chromosomes at a similar rate (Supplementary Figure 5C). On the other hand, we found fewer chromosome segregating cells at 60 min post release into mitosis when TRAIP was absent (Supplementary Figure 5D). This suggests that TRAIP is indeed regulating the timing of mitotic progression independently of its S-phase role. Therefore, to determine whether any functions of TRAIP in mitosis could contribute to the increased DNA damage response seen in the next S-phase, TRAIP was degraded specifically before mitosis and its effect on the next cell cycle arrest was explored. Cells were arrested in G2 and treated with auxin (IAA) for 1 hour prior to release into mitosis. Approximately 2 hours following release, when cells were in G1 phase of the cell cycle, auxin (IAA) was optionally washed off to allow protein re-expression before entry into the next S-phase. Timepoints were taken 24 hours after release into mitosis. Using this experimental set-up, cell cycle accumulation in G2 was only seen when cells progressed through both mitosis and S-phase without TRAIP. No such accumulation was seen in those samples where TRAIP was allowed to re-express prior to S-phase entry (Figure 3C). We therefore conclude that the essential function of TRAIP is executed during S-phase.

### TRAIP depletion leads to generation of DNA damage at transcription start sites (TSS)

Given the essential requirement of TRAIP during S-phase progression, we explored whether auxin (IAA) treatment had any effect on global DNA replication. Cells treated with and without auxin (IAA) for 24 hours were pulsed with the thymidine analogues EdU and BrdU for 1 hour, and analogue incorporation was analysed using immunofluorescence and FACS respectively (Supplementary Figure 6A,B). Whilst overall lower proportions of cells were detected incorporating EdU/BrdU, no differences in the average amount incorporated per positive cell were found (Supplementary Figure 6A,B). Based on our previous observations, it is likely that the lower proportions of cells in S-phase are due to the G2 cell cycle arrest and overall fewer cycling cells. TRAIP has also been previously implicated in replisome disassembly in model organisms during mitosis and stressed conditions ^12,17,20–23^. We therefore wondered whether TRAIP was needed for global replisome disassembly during S-phase. The levels of chromatin-bound replicative helicase (MCM2-7) were assessed using flow cytometry following extraction of soluble proteins from permeabilised cells. No differences could be detected in either the loading or unloading of MCM2-7 during the otherwise unperturbed cell cycle (Supplementary Figure 6C). Despite these results, the S-phase progression upon TRAIP degradation is not without problems. To investigate this further, cells progressing through S-phase with or without TRAIP were arrested in G2 with RO-3306 and pulsed with EdU to detect any late cell cycle DNA synthesis. When cells progressed through S-phase without TRAIP, our results suggest that they struggled to complete DNA synthesis on time as EdU incorporation could be detected in late G2/early mitosis (Supplementary Figure 7A).

In order to gain insight into the mechanistic requirements for TRAIP during S-phase, we used γ-H2AX ChIP-Seq to map genome-wide the location of the accumulating DNA damage. We first confirmed that the previously observed DNA damage accumulation was indeed specific to TRAIP’s function during S-phase (Supplementary Figure 7B). Cells were then prepared for sequencing at the 16-hour timepoint (late S-phase) as determined to display the maximal accumulation of DNA damage (Supplementary Figure 7B). H2AX chromatin immunoprecipitation was used for sample normalisation and we performed two independent repeats of this experiment. As per our previous data, the ChIP-Seq detected overall increases to the γ-H2AX signal detected following auxin (IAA) treatment in both repeats (example in Figure 4A). We first checked whether the position of γ-H2AX signal correlated with known DNA replication features: replication initiation sites and replication termination zones (both mapped in HCT116 cells by alternative techniques by^33^ and^32^) and common fragile sites (CFS)^42^. However, no correlation could be found in either of the repeats (Figure 4B and Supplementary Figure 8A). Subsequently we performed a peak calling analysis to determine hotspots of DNA damage in the two repeats. We determined 1799 γ-H2AX signal peaks in the first experiment and 5984 in the second. 545 γ-H2AX signal peaks were common between both experiments (example of a hotspot common peak is presented in Figure 4A). Interestingly, when we analysed the location of these 545 γ-H2AX hotspots, we found that they were largely associated with RNA Pol II transcripts. Indeed, we found that 95.77% of these peaks were present on genes, and 88.24% present specifically around the transcription start sites (TSS) +/- 2kb (Figure 4C). We could also see a similar correlation with TSS for all γH2AX peaks called in both experiments, not only the common hotspots (data not shown). Guided by this finding, we analysed the positions of all TRAIP-degradation induced γH2AX signals in both repeats in relation to all transcribed genes. Importantly, we found a general increase of γ-H2AX signal at transcription start sites, even when the damage levels were not increased enough for stringent conserved peak calling analysis (Figure 4D and Supplementary Figure 8B). Hence, we focused on understanding what was special about the TSS of the genes exhibiting the hotspots of DNA damage upon TRAIP degradation. Firstly, we excluded an enrichment for genes involved in any particular cellular pathway through Go-term analysis (Supplementary Table 1). Next, we characterised replication dynamics across these sites using previously published replication data for HCT116 ^33^, to determine the replication timing associated with these sites and we found that hotspot TSS sites are preferentially replicated in early S-phase (Supplementary Figure 8C). We then identified the closest replication origin, as assigned by *Koyanagi et al*. to the γ-H2AX peak^32^, determining also whether the replication fork will reach the TSS in a codirectional or head-to-head orientation with the RNA Pol II. We found that γ-H2AX peaks are close to their nearest origin, on average only 29 kb away (Figure 4E). In comparison, TSS of genes that do not show an increase in γ-H2AX levels following TRAIP depletion (top 450 genes Supplementary Fig 8B) are 41 kb away from their nearest origin (Figure 4E). Moreover, γ-H2AX peaks occur preferentially around the TSS of the first transcribed gene encountered by the replication fork (75% of the cases), with same frequencies of replication fork and gene orientation being codirectional or head-to-head (Figure 4F). In comparison, TSS of the genes with lowest changes in γ-H2AX levels tend to occur less frequently on the first transcribed gene encountered by the replication fork (65% of the cases, Figure 4F). We then overlayed this information with previously published datasets for precision run-on sequencing (PRO-Seq), tracking strand specific nascent transcription activity ^29^, as well as RNA Pol II chromatin immunoprecipitation sequencing (ChIP-Seq)^30^ in HCT116 cells. Interestingly, when we analysed the PRO-Seq signal at the hotspot TSS, we found that 98.25% of all the genes with a γ-H2AX hotspot presented clear levels of bi-directional transcription at their TSS, independently of the reciprocal orientation, either because of bidirectional promoters or TSS-associated antisense transcription. We measured therefore the extent of the antisense transcription by calculating the ratio between the levels of the antisense transcription and the sense transcription. To do this, we used the strand specific PRO-Seq data, and determined for each gene the amount of antisense transcription occurring in the region −1000bp -> TSS on the opposite strand of the gene. This was then divided by the amount of sense transcription TSS -> +1000bp. We found that, compared to all the rest of the transcribed genes, those with a γ-H2AX hotspot peak present higher levels of antisense transcription at their TSS (Figure 4G). We also did the same analysis with total RNA Pol II ChIP-Seq data ^30^ finding the same result (Figure 4G, Supplementary Figure 8D). In comparison, genes with the lowest changes of γ-H2AX levels at their TSS present lower levels of antisense transcription both by PRO-Seq and ChIP-Seq of RNA Pol II (Figure 4G). Altogether, our data indicate that TSS genomic regions are particularly challenging to replicate in the absence of TRAIP. Moreover, although γ-H2AX signal is generally increased at the majority of TSS, it is the TSS of the first transcribed gene with high levels of antisense to sense transcription that DNA damage will reproducibly arise at.

**Figure 4.**
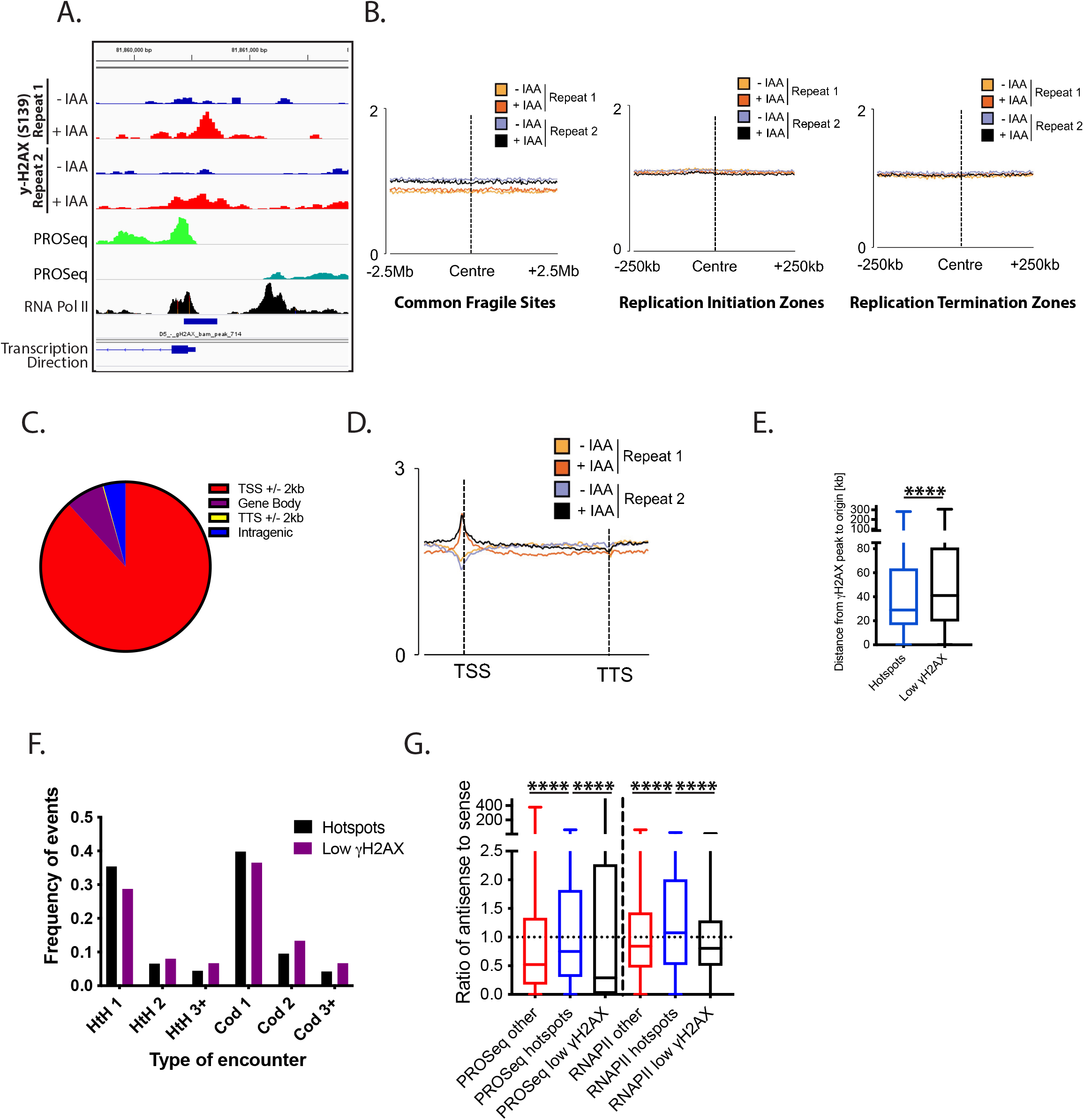
Absence of TRAIP during S-phase leads to generation of DNA damage at transcription start sites (TSS). **(A)** An example of TRAIP-degradation induced γH2AX peak over two independent repeats of γH2AX ChIP-seq experiments. Transcription direction and position of accumulation of RNA Pol II ^30^ and of PRO-Seq ^29^ around the promoter are also presented. **(B)** There is no correlation between γH2AX ChIP signal and replication initiation or termination zones nor common fragile sites. γH2AX ChIP signal from two independent experiments was compared with positions of replication initiation and termination zones as mapped by ^33^, or common fragile sites as mapped by^42^. Each graph is centred around the centre of initiation/termination/fragile site. **(C)** Graphical representation of the position of the called conserved γH2AX peaks (damage hotspots), with 88.24% of γH2AX peaks in proximity of TSS. **(D)** Average binned metagene profile for γH2AX levels normalised to H2AX at transcribed genes in HCT116 cells, from TSS to transcription termination sites for the two repeat experiments. **(E)** Distance between the γH2AX peaks at TSS at hotspots and the TSS of genes with the lowest fold change in γH2AX levels and origins of replication identified in *Koyanagi et al*., 2021. The positions of TSS with called common γH2AX peaks is significantly closer to origin of replication than the distance between TSS and origin of replication in genes that do not show increased γH2AX signal upon TRAIP degradation. Box whiskers plots with line at the median; Mann-Whitney t-test; **** = p-value < 0.0001. **(F)** Called γH2AX peaks are located preferentially at TSS of first gene encountered by replication forks emerging from closest replication origins either in head-to-head or codirectional mode. The proportion of 1^st^, 2^nd^ and further on (3+) genes are presented. Genes with lowest fold changes in γH2AX levels are less preferentially the first gene encountered by replication forks emerging from the closest replication origin. Chi-square p-value < 0.05. **(G)** Quantification of the ratio between antisense- and sensetranscription at TSS +/- 1kb of hotspots, all other transcribed genes and transcribed genes with low levels of γH2AX increase following TRAIP depletion. PRO-Seq strand specific signal^29^ and RNA Pol II chromatin immuno-precipitation sequencing ^30^ was quantified −1000 bp to TSS for antisense transcription, and TSS to 1000 bp for the sense transcription. Box whiskers plots with line at the median; Mann-Whitney t-test; **** = p-value < 0.0001.

### TRAIP is required to resolve DNA replication - transcription conflicts on chromatin

TRAIP was shown previously to be able to remove a DNA-protein crosslink (DPC) barrier in front of the replisome ^17^. As we observed that DNA damage was preferentially located at transcription start sites following TRAIP degradation, we hypothesised that the RNA polymerase accumulated at these sites could present a barrier for the passing replisome. To test this, we decided to inhibit loading of the RNA Pol II at TSS using the TFIIH inhibitor triptolide ^30^. Cells were arrested in G1 where TRAIP was degraded before being released into S-phase. Upon S-phase entry, cells were exposed to triptolide for 90 minutes, as it has been shown previously that short term treatment with triptolide does not affect DNA replication progression ^43^ (Supplementary Figure 9). Strikingly, all of the DNA damage observed upon TRAIP degradation could be rescued by inhibiting RNA Pol II recruitment to the TSS by triptolide treatment (Figure 5A). We also treated cells in an analogous way with another transcription inhibitor DRB. DRB is a CDK9 inhibitor which leads to transcription inhibition through accumulation of RNA Pol II at the TSS, inhibiting its progression through the gene body ^44^. We could observe that DRB treatment alone created an increased level of γ-H2AX and 53BP1 foci, similar to that of TRAIP degradation. DRB treatment did not lower the proportion of cells displaying increased number of γ-H2AX and 53BP1 foci upon TRAIP degradation, but the effect of DRB treatment was also not additive with TRAIP degradation, suggesting that they act in the same pathway (Figure 5B). Altogether, these data suggest that the mechanism by which TRAIP is essential during S-phase is dependent on the presence of RNA Pol II at the TSS.

**Figure 5.**
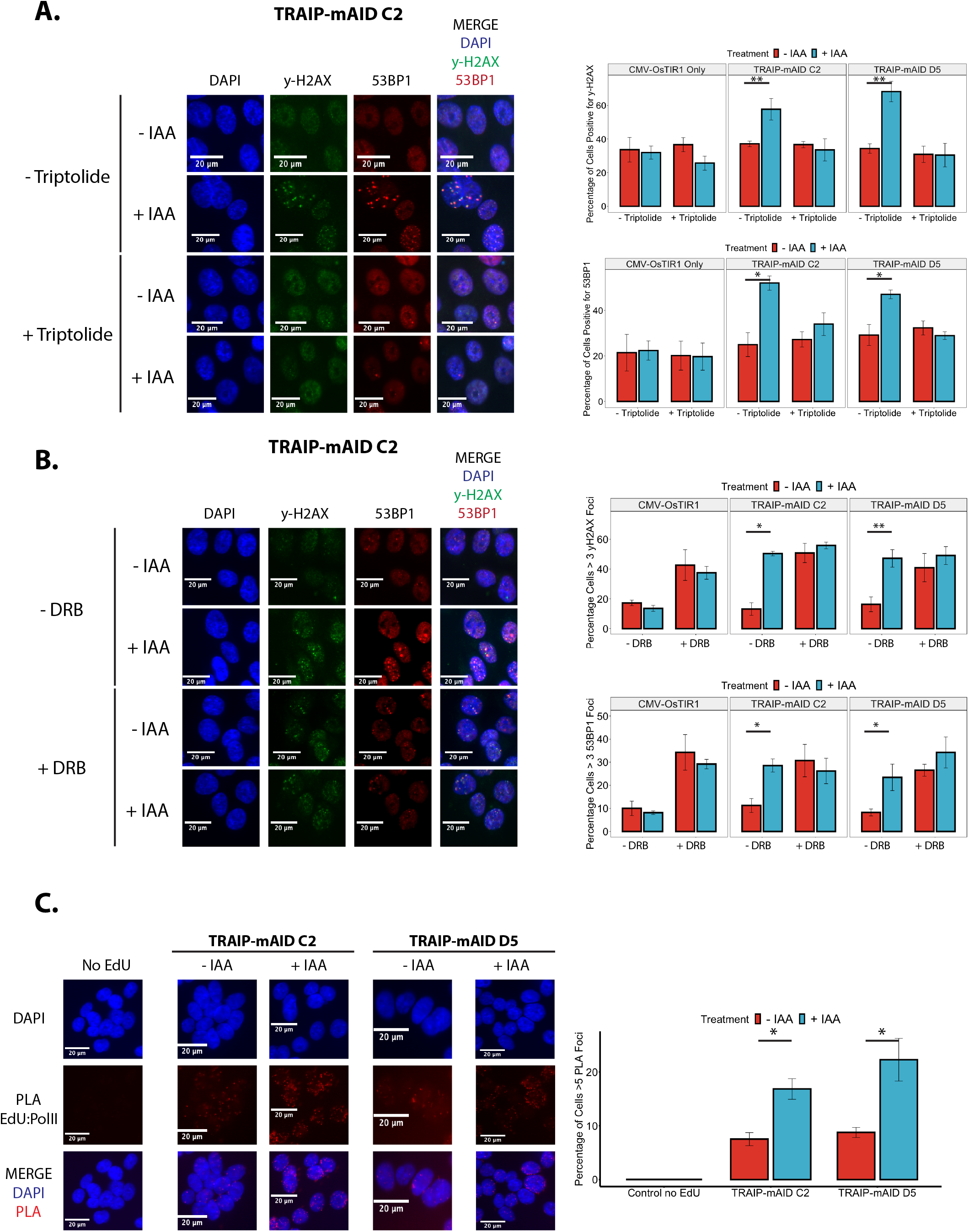
DNA damage induced in S-phase by TRAIP degradation depends on presence of transcription machinery. **(A)** Inhibition of RNA Pol II loading onto chromatin with triptolide rescues TRAIP-degradation-induced DNA damage. TRAIP-mAID cells were arrested in G1 stage of the cell cycle and TRAIP degraded before release of cells into S-phase. Upon S-phase entry cells were exposed to triptolide for 90 min. γH2AX and 53BP1 foci induced were visualised by immunofluorescence and quantified. Example pictures (left) and quantification of 3 independent experiments (right) is presented (mean -/+ SEM).). Differences between conditions was identified using t.tests (yH2AX: TRAIP-mAID C2 - Triptolide: p = 0.00622; TRAIP-mAID D5 - Triptolide: p = 0.00798. 53BP1: TRAIP-mAID C2 - Triptolide: p = 0.0424; TRAIP-mAID D5 - Triptolide: p = 0.0474). **(B)** TRAIP degradation and inhibition of progression of RNA Pol II are epistatic. Experiment as in (A) but cells were treated for 90 min with DRB rather than Triptolide. Differences between conditions was identified using t.tests (yH2AX: TRAIP-mAID C2 -Triptolide: p = 0.0146; TRAIP-mAID D5 - Triptolide: p = 0.00738. 53BP1: TRAIP-mAID C2 - Triptolide: p = 0.0243; TRAIP-mAID D5 - Triptolide: p = 0.0259). **(C)** Lack of TRAIP in S-phase leads to an increased number of transcription machinery collisions with nascent replicated DNA. TRAIP-mAID cells were arrested in G1 stage of the cell cycle and TRAIP degraded before release of cells into S-phase. In early S-phase EdU was added for 20 min and proximity between incorporated EdU and RNA Pol II visualised by PLA. Example pictures (left) and quantification of cells with over 5 PLA foci over 3 independent experiments (right) is presented (mean -/+ SEM). Pairwise hypothesis testing was carried out using t.tests (TRAIP-mAID C2: p = 0.02; TRAIP-mAID D5: p = 0.0442).

Conflicts or collisions between the replicative helicase and transcription machineries are known to be major sources of genomic instability in otherwise unperturbed cells; through both interactions between the protein complexes themselves, or formation of DNA:RNA hybrids (R-loops)^45^. Given the requirements of Pol II bound DNA for the accumulation of DNA damage after auxin (IAA) treatment, it is likely that TRAIP is required for the resolution of conflicts between DNA replication and transcription. To test this, we used proximity ligation assays (PLA) to explore any differences in the proximity of active transcription and nascent DNA. Cells were arrested in G1 where TRAIP was degraded before being released into S-phase.

Approximately 12 and a half hours following release when cells were in very early S-phase (known to have high levels of both transcription and replication and when our γH2AX hotspots are replicated), the thymidine analogue EdU was added for 20 minutes. Cells were then harvested and PLA assays used to detect interactions between active transcription (phosphorylated RNA Pol II) and nascent replicated DNA (Figure 5C). Intriguingly, we observed an increase in the amount of active Pol II present in proximity to nascent DNA following TRAIP degradation, suggesting that indeed TRAIP is important for resolving replication-transcription conflicts on chromatin and maintaining fork progression.

### TRAIP depletion does not lead to DNA damage during S-phase in absence of transcription

To explore further the importance of TRAIP for the regulation of replication-transcription collisions we next turned to the *Xenopus laevis egg* extract model system, which can support robust DNA replication activity in absence of any transcription. During early embryogenesis in *Xenopus* embryos 12 cleavage cell cycles are achieved without transcription and only restricted protein translation. Most required factors for DNA replication and cell division are accumulated in the egg. The significant level of gene transcription is induced in the embryo only at the stage of midblastula transition. *As Xenopus egg* extract is derived from *Xenopus eggs* and the DNA substrate replicated is a demembranated *Xenopus* sperm, the replication observed resembles embryonic replication during early cleavage divisions in absence of transcription.

Using this system, we immunodepleted TRAIP from *Xenopus egg* extract to less than 10% of its original quantity (Figure 6A) and could observe that although a lack of TRAIP in the extract did not affect its ability to synthesise DNA during S-phase, as previously reported ^22^, it did inhibit mitotic unloading of post-termination replisomes, which is a known function of TRAIP in this system (Supplementary Figure 10B). This verified that our immunodepleted extract was indeed devoid of TRAIP’s activity. Given our findings in the mammalian system, we next examined whether we could observe any signs of creation of DNA damage or checkpoint activation during S-phase without TRAIP in *Xenopus egg* extract system. Interestingly, we could detect no increase of γ-H2AX signal on chromatin, nor phosphorylated Chk1 in the nucleoplasm in the absence of TRAIP (Figure 6B and C). This further supports our hypothesis that TRAIP works in S-phase to protect genome stability by resolving conflicts between replication and transcription machineries; in the absence of transcription, such DNA damage does not occur.

**Figure 6.**
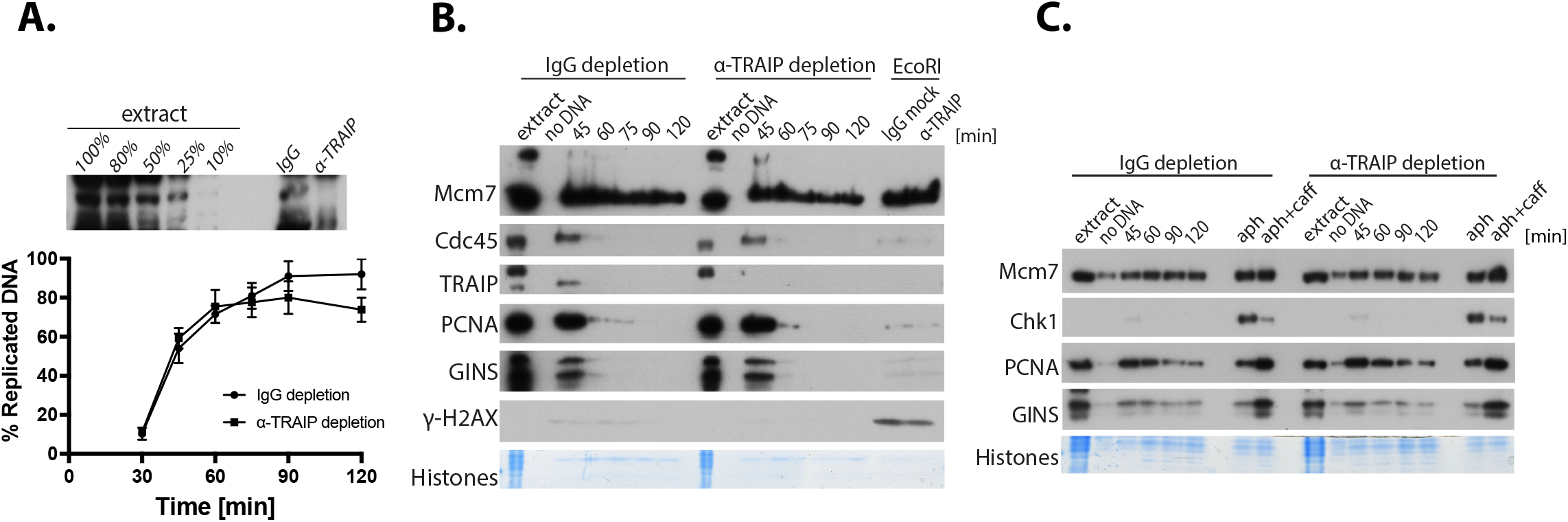
Depletion of TRAIP does not induce S-phase checkpoint activity in *Xenopus laevis* egg extract. **(A)** TRAIP is not essential for S-phase progression in *Xenopus laevis egg* extract. TRAIP was immunodepleted from *Xenopus egg* extract to less than 10% as judged by immunoblotting of depleted extract (top). The ability of TRAIP-depleted extract to synthesise DNA was established by measuring α^32^P-dATP incorporation into newly synthesised DNA and compared with replication of non-specific IgG-depleted extract. **(B)** No increase in H2AX phosphorylation during DNA replication is detected upon TRAIP depletion. DNA replication reaction was established in non-specific IgG- and TRAIP-depleted egg extracts and chromatin isolated at indicated timepoints during replication reaction. Chromatin samples were analysed by western blotting with indicated antibodies. A “no DNA” sample shows chromatin specificity of the signal. Histones stained with Coomassie serve as a loading control. Samples of both extracts treated with restriction enzyme EcoRI to induce double strand breaks (isolated at 60 min of reaction) serve as positive control for γH2AX signal. **(C)** Active Chk1 is not detected during DNA replication in TRAIP-depleted *Xenopus egg* extract. DNA replication was established as in (B) but instead of chromatin whole nuclei containing nucleoplasm were isolated to measure level of active phosphorylated Chk1. Samples of both extracts treated with the DNA Polymerase inhibitor aphidicolin serve as positive controls for Chk1 activation, while samples treated with aphidicolin and caffeine indicate ATM/ATR dependence of these signals.

**Figure 7.**
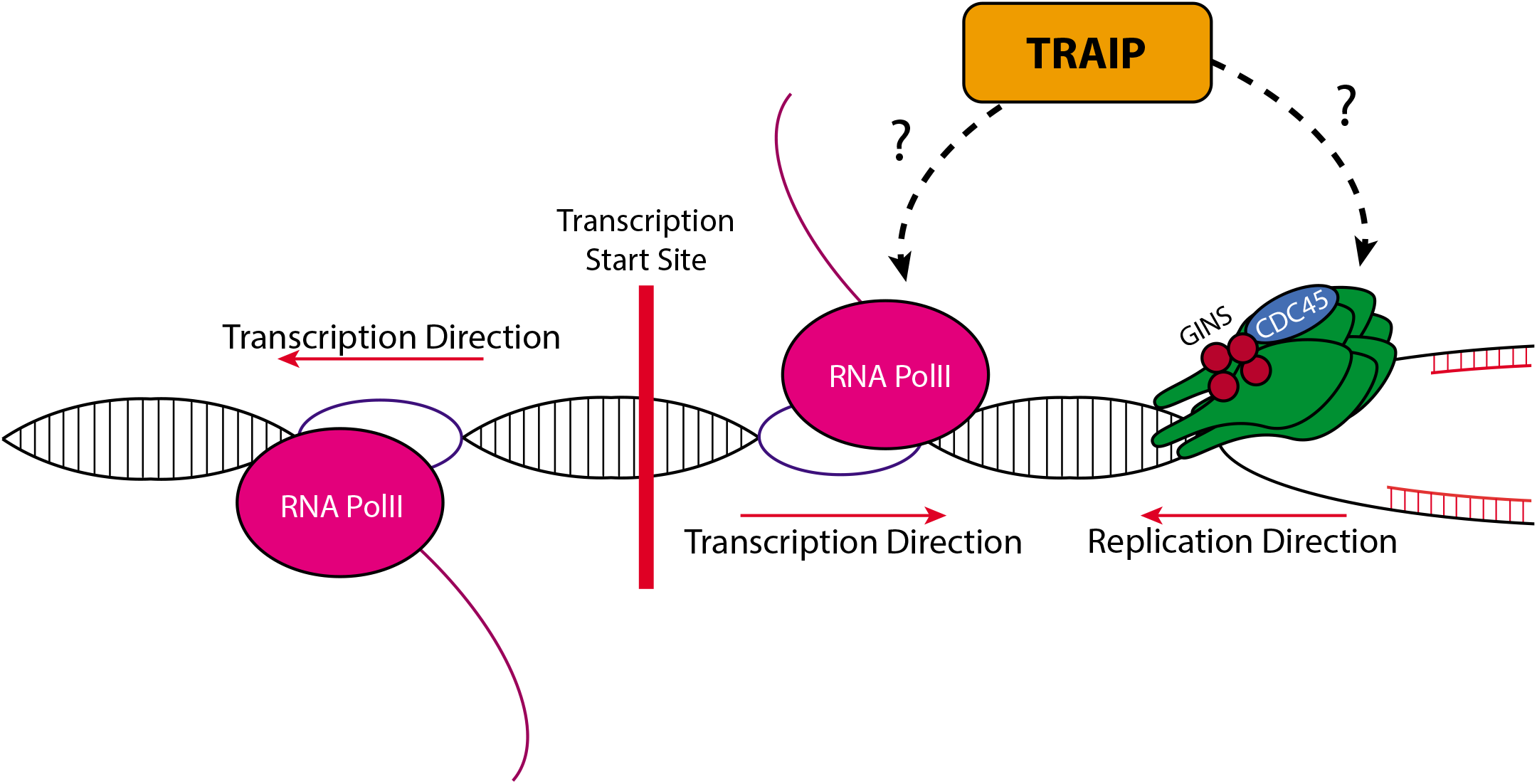
Model of TRAIP activity during S-phase.

## Discussion

Over the years, much evidence has accumulated indicating that TRAIP ubiquitin ligase plays wide-ranging roles in the maintenance of genomic integrity. TRAIP-depleted cells were shown previously to display a diverse spectrum of phenotypes: gross chromosomal re-arrangements, problems with DNA damage response activation, micronuclei, accumulation in G2 stage of the cell cycle ^5–7^. The specific functions of TRAIP were studied upon specific stimuli e.g. induction of double strand breaks, induction of inter-strand crosslinks, stimulation of MiDAS or inhibition of replicative polymerases etc.^6,12,18,21^. However, the primary consequence of TRAIP depletion, the essential function of TRAIP for cell viability in unperturbed conditions, was not known. Here, we have taken advantage of the degron system to rapidly degrade TRAIP within cells upon auxin (IAA) addition (TRAIP-mAID), to dissect the importance of various functions of TRAIP in different stages of the cell cycle. Firstly, we have shown that upon the degradation of TRAIP, cells show signs of spontaneous DNA damage, leading to ATR checkpoint activation, G2 cell cycle arrest and withdrawal from the cell cycle into senescence. TRAIP-deficient human cells have been shown to proliferate slowly even in the absence of exogenous DNA damage and the reduction of cellular proliferation during development is a likely cause of microcephalic dwarfism observed in patients with TRAIP mutations ^5,46^. Here, for the first time we have determined the mechanism of inhibition of cell proliferation in TRAIP-depleted cells. Interestingly, despite TRAIPs function to unload rogue replisomes during mitosis ^20–22^, the promotion of MiDAS ^20–22,47^, or involvement in the spindle assembly checkpoint^9,24^, any insults to genome stability generated by cells progressing through mitosis without TRAIP are not responsible for the perturbed cell proliferation and G2 arrest. Instead, we find that the functions of TRAIP during S-phase are of utmost importance for cell viability.

TRAIP has been shown previously to be important for the cellular response to DNA replication stress upon different types of insults ^5–7^. It has also been suggested to be a master regulator of ICL repair during DNA replication, as its activity promotes two alternative pathways of ICL repair in *Xenopus laevis egg* extract^12^. However, patients with TRAIP mutations display dwarfism rather than classic Fanconi anaemia clinical outcome, which is characteristic for mutations within ICL repair factors ^5,48^. Instead, our results indicate, that without exogenous sources of replication stress, the major endogenous source of replication fork impediment that requires TRAIP activity is in encountering transcription machinery. In the absence of TRAIP during S-phase in human cells, we observe increased levels of persisting DNA replication-transcription collisions. We can also detect enrichment of the DNA damage response at TSS sites, where replication forks likely collide with RNA Pol II. Importantly, this damage can be completely rescued by the temporary removal of RNA Pol II from chromatin. Moreover, in *Xenopus egg* extract model system where efficient DNA replication can be established without the presence of concurrent transcription, TRAIP is not essential for the completion of S-phase, nor can we detect any signs of DNA damage or checkpoint activation.

In human cells, coexistence of DNA transcription and DNA replication is a well-established potential source of endogenous replication stress, that is often exacerbated by oncogenic deregulations stimulating transcription, whilst simultaneously promoting premature entry into S-phase, leading to a higher probability of interference between the two processes ^43 49^. The role of TRAIP in resolution of such conflicts is consistent with the observation that TRAIP-deficient cells show fork asymmetry in the DNA fibre assays, suggesting site specific stalling of one of the forks emanating from the same origin of replication ^6^.

What is the mechanism by which TRAIP may promote resolution of replication-transcription collisions? Our analysis of the hotspots of DNA damage created in the absence of TRAIP suggests that the problems most often arise when the newly established replication fork encounters the first gene promoter where RNA Pol II has accumulated. Moreover, the hotspot sites represent particularly crowded sites, with high levels of transcription moving in both directions from the promoter. Despite the orientation of gene transcription being equally often in head-to-head or codirectional with the direction of progression of the replication fork, the equal levels of sense to antisense transcription at hotspot TSS suggest that the fork will more often encounter the transcription machinery in a head-to-head orientation. These bi-directional transcription units may therefore represent a particularly difficult impediment for replication fork progression. Indeed, sites with high levels of antisense transcription have already been identified as hotspots for transcription-replication interactions and for G2/M DNA synthesis ^28,50^. Previous research has also identified that TSS-associated transcription is stimulated by the presence of R-loops accumulating at the TSS ^51^ and the propensity to form R-loops is a feature of transcription replication interaction hotspots ^50^. However, when we assessed R-loops levels using the R-loop specific antibody S9.6 by immunofluorescence, we did not identify an increase in cells depleted of TRAIP, indicating that TRAIP depletion is not leading to a global increase in R-loops levels in cells (Supplementary Figure 9B).

The transcription machinery encountered by the replication fork very much resembles a DNA-protein barrier and TRAIP has been shown previously to facilitate replisome bypass of DNA protein crosslinks (protein covalently crosslinked to DNA by aldehydes or chemotherapeutics) ^17^. In this situation, TRAIP stimulates the bypass of the DPC by the replication machinery and also ubiquitylates DPC to stimulate its degradation by SPRTN and the proteasome ^17,52^. It is therefore likely that TRAIP could act similarly when encountering tightly associated RNA Pol II. It is well established that removal of RNA Pol II from chromatin in response to DNA damage is driven by ubiquitylation of RNA Pol II and subsequent unfolding and unloading by p97 segregase^53^. Further research with site specific collision of both machineries will be needed to establish the precise molecular chain of events.

Finally, downregulation or immunodepletion of TRAIP in human cells and *Xenopus egg* extract system was reported previously to lead to chromosomal instability and rearrangements, which are a hallmark of cancer development^6,7,22^. Notably, however, TRAIP deficient patients were not reported to characterise with cancer predisposition, similarly to other Seckel syndrome patients^5,46^. It is possible therefore that the G2 arrest and senescence we observe upon degradation of TRAIP is not compatible with increased proliferation and tumour development.

Inhibition of TRAIP could therefore be significantly detrimental to cancerous cells. In an analogous manner, inhibitors of ATR are currently in development as promising cancer treatments for tumours lacking certain DNA damage response pathways ^54^. Indeed, we find that a combination of ATRi and TRAIP degradation leads to an additive increase in cells displaying elevated levels of γ-H2AX and 53BP1 foci (Figure 8). This result suggests that TRAIP inhibition alone could be a promising target for future cancer therapy, to cease the growth of cancerous cells, but could also work well in combination with recently developed ATR inhibitors ^54^.

**Figure 8.**
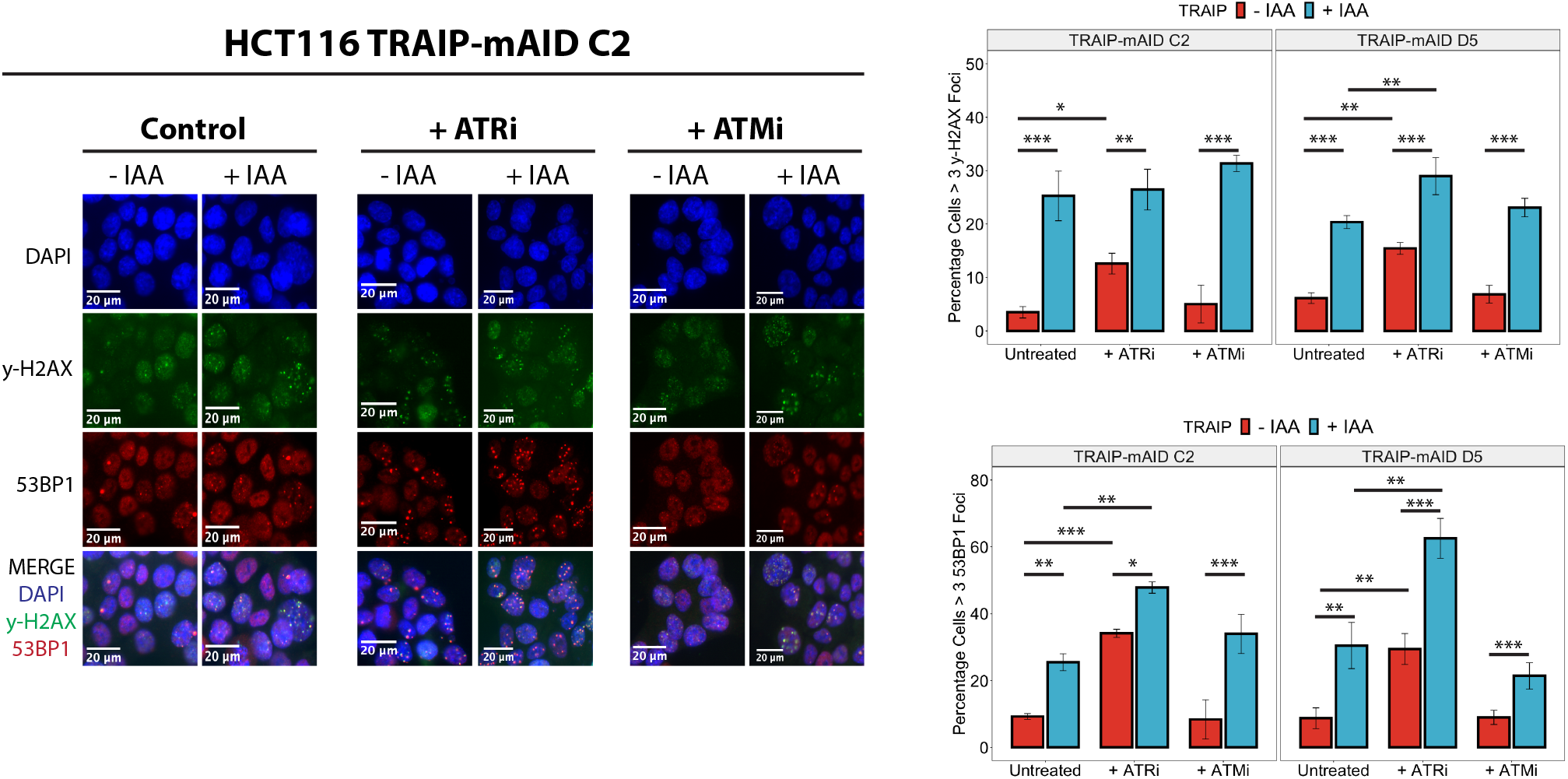
TRAIP degradation sensitises cells to ATR inhibition. TRAIP-mAID cells were treated optionally with auxin (IAA) for 24 h and indicated checkpoint kinase inhibitor. γH2AX and 53BP1 foci induced were visualised by immunofluorescence and quantified. Example pictures (left) and quantification of 3 independent experiments (right) is presented Quantification shown as mean percentage of cells +/- SEM.

## Supporting information

combined supplementary figures

## Acknowledgments

This work was supported by the BBSRC funded MIBTP studentship and JSPS Summer programme for S.S., and Wellcome Trust Investigator Award (215510/Z/19/Z) for A.G. University of Birmingham, BBSRC (BB/S016155/1) and Cancer Research UK (C17422/A25154) to M.S.

