## Supplementary material for "Ubiquitin ligase TRAIP plays an essential role during the S-phase of unperturbed cell cycle in the resolution of DNA replication – transcription conflicts": combined supplementary figures

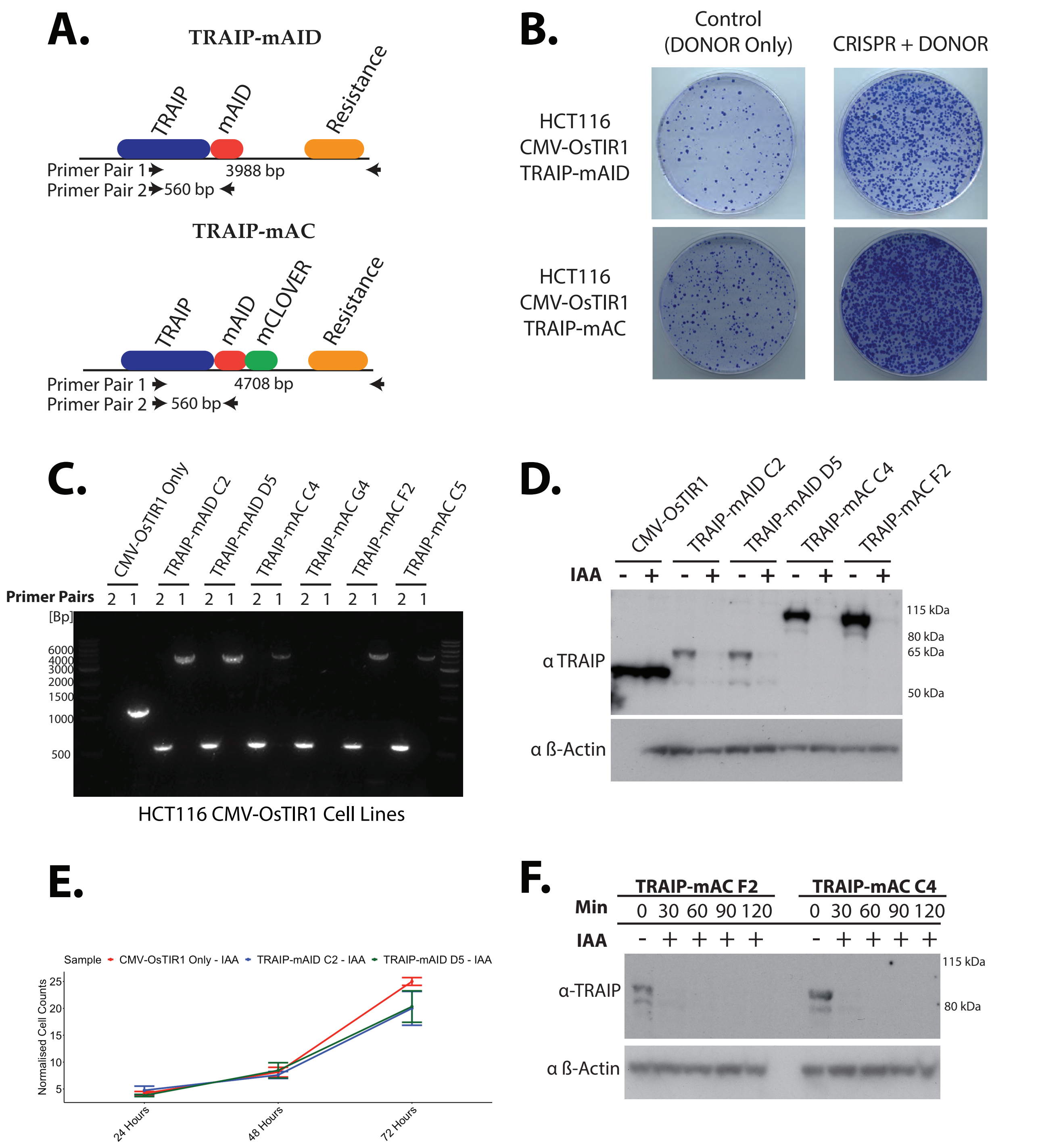


**Supplementary Figure 1. Creation and validation of TRAIP degrons in HCT116 cells. (A)** Design of TRAIP tagged with auxin inducible degron (mAID) or mClover tagged degron (mAC). Primer pairs used for verification of correct genomic incorporation is presented below the diagrams. **(B)** Examples of cell colonies obtained after guide RNAs transfection stained with crystal violet. **(C)** Validation of biallelic tagging of TRAIP in different clones using primer pairs indicated in (A). **(D)** Validation of biallelic tagging of TRAIP by western blotting with TRAIP antibodies. **(E)** Growth curves of control (CMV-OsTIR1) and two clones of TRAIP-mAID cells ($n = 3$). Hypothesis testing was carried out using one-way ANOVA for each individual timepoint (24 h: $Df = 2$, $F = 0.892$, $p = 0.458$. 48 h: $Df = 2$, $F = 0.18$, $p = 0.839$. 72 h: $Df = 2$, $F = 1.236$, $p = 0.355$). **(F)** Degradation of TRAIP-mAC within 30 min of auxin addition.

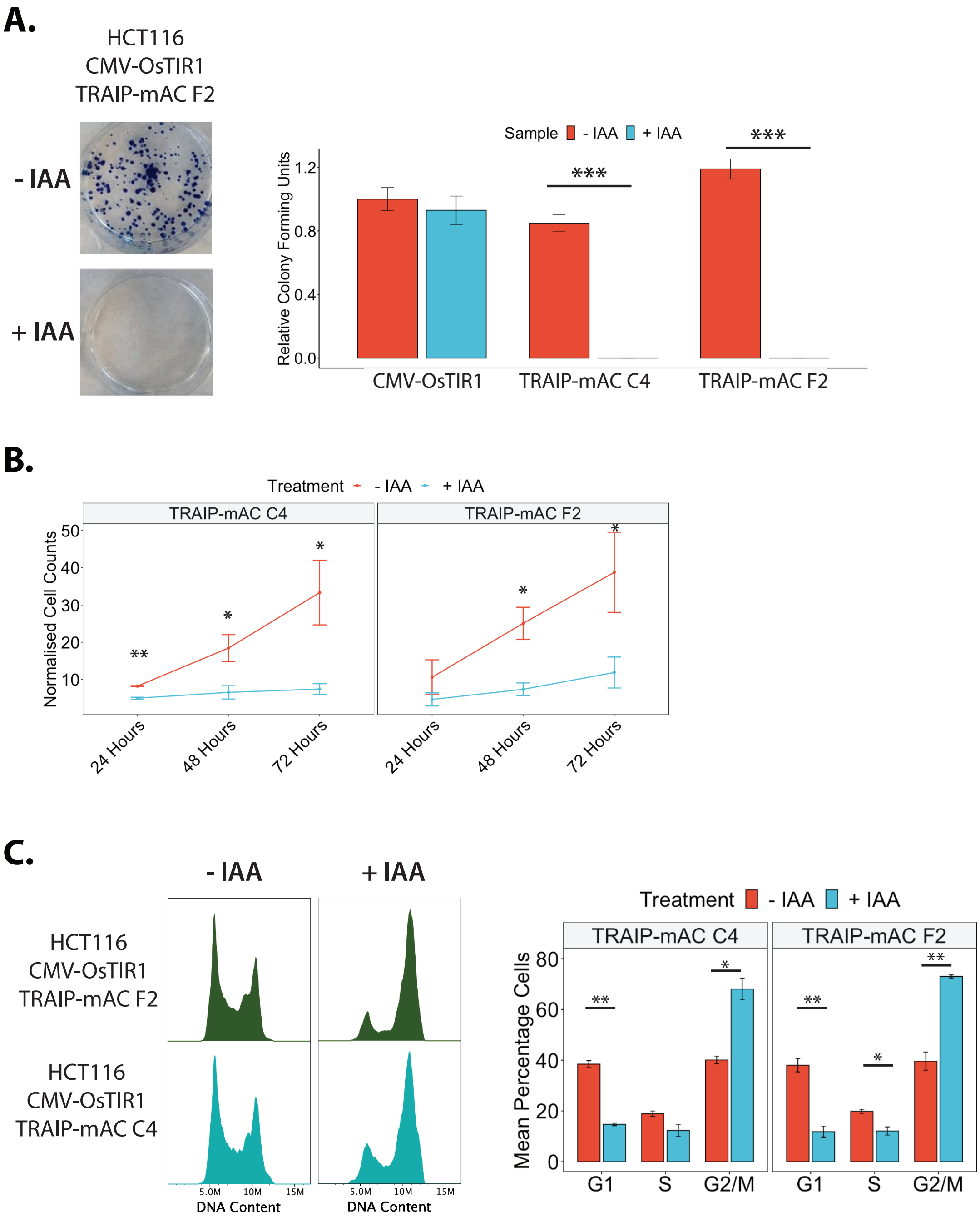


**Supplementary Figure 2. Phenotypes of inhibited proliferation and cell cycle arrest upon TRAIP degradation are reproduced in TRAIP-mAC cell lines. (A)** Example of colony assay after auxin treatment. Cells were fixed and stained with methylene blue (left). Quantification of colony forming assay in two independent TRAIP-mAC clones n=3 (right). Statistical analyses was carried out using T.tests. Significant differences were identified between TRAIP-mAC C4 (p = 0.000161) and TRAIP-mAID F2 (p = 0.000159). **(B)** Growth curves of TRAIP-mAC cells upon 24, 48 and 72 h of auxin treatment. Cells were counted at every timepoint and normalised to the seeding densities n=3. Statistical significance was calculated using T.tests for TRAIP-mAC C4 (24 hours: p= 0.002158; 48 hours: p= 0.03; 72 hours: p= 0.04613) and TRAIP-mAC F2 (48 hours: p = 0.0399; 72 hours: p = 0.03326). **(C)** TRAIPmAC clones arrest at G2/M stage of the cell cycle. examples of FACS cell cycle profiles of two TRAIP-mAC clones upon 24 h of optional auxin treatment (left). Quantification of cell cycle stages in TRAIP-mAC clones after 24 h auxin treatment n=3 (right). Statistical analysis (t.tests) - G1: TRAIP-mAID C4 (p = 0.00122), and TRAIP-mAID F2 (p = 0.00175); G2/M - TRAIP-mAC C4 (p = 0.0153), TRAIP-mAC F2 (p = 0.00978).

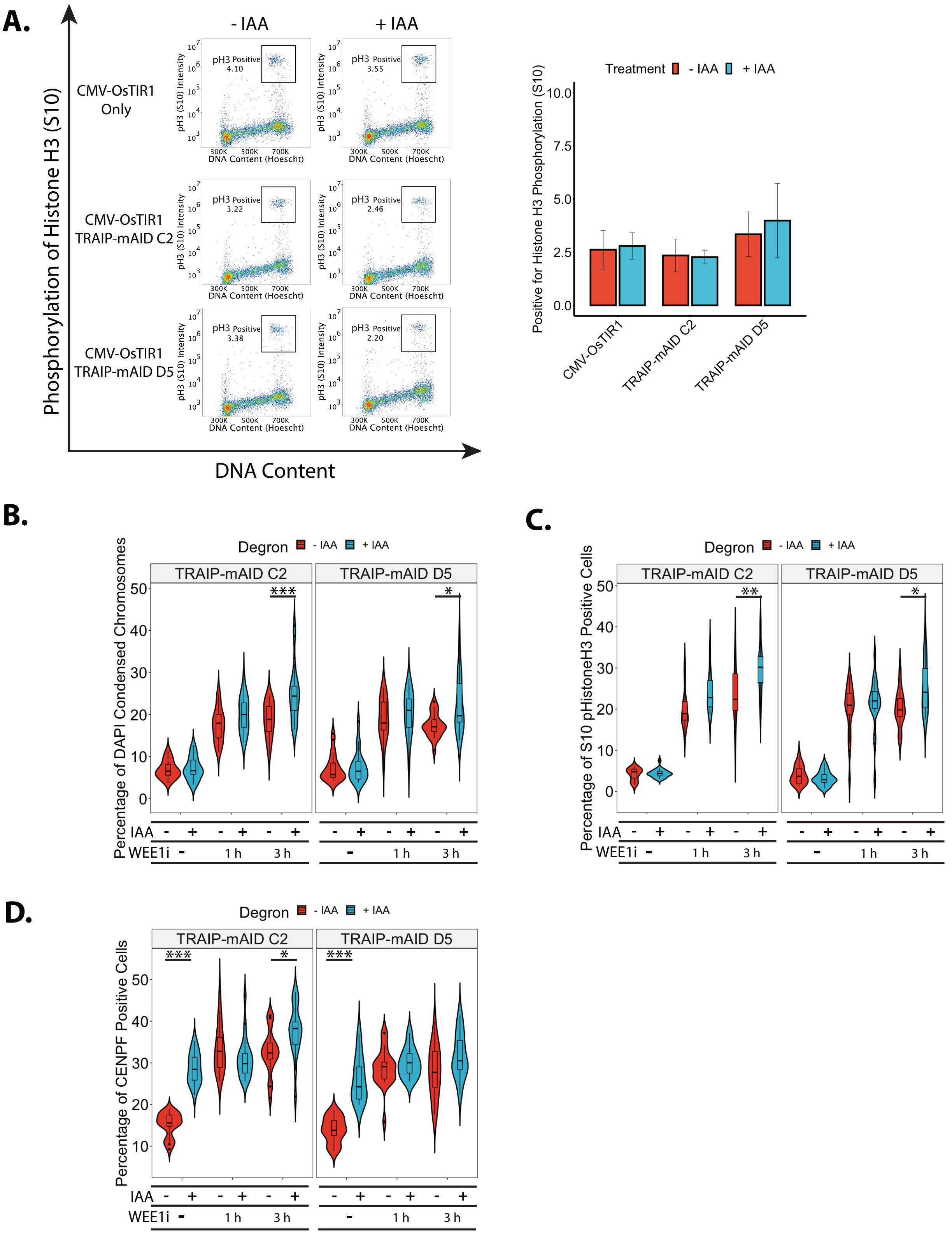


**Supplementary Figure 3. TRAIP depleted cells accumulate in G2 stage of the cell cycle. (A)** Degron cell lines were treated with auxin for 24 h and stained for S10 phosphorylation on histone H3 (pH3). Example FACS plots with pH3 staining against DNA content. Mitotic cells are selected in the black box (left). Quantification of the percentage of cells in mitosis (pH3 positive) upon 24 h of TRAIP degradation n=3 (right). Statistical analysis revealed no differences in the data. **(B)** Quantification of the percentage of cells found to be in mitosis after 24 h of auxin treatmentby quantification of cells with condensed chromosomes after DAPI staining and fluorecent microscopy (mitotic index). Significance (ANOVA and pairwise testing) is summarised on the plot: TRAIP-mAID C2 ($p = < 0.001$) and TRAIP-mAID D5 ($p = 0.0162$). **(C)** Quantification of H3S10 phosphorylation by fluorescent microscopy in TRAIP degron cell upon 24 h of auxin treatment and optional treatment with WEE1 inhibitor MK-1775. Significance (ANOVA and pairwise testing) is summarised on the plot: TRAIP-mAID C2: $p = 0.00260$; TRAIP-mAID D5: $p = 0.0170$. **(D)** Quantification of CENPF positive cells upon treatment as in (C). Significance (ANOVA and pairwise comparison) is summarised on the plot: TRAIP-mAID C2 Untreated ($p = < 0.001$), TRAIP-mAID C2 + Wee1i 3 hours ($p = 0.0294$), and TRAIP-mAID D5 untreated ($p = < 0.001$).

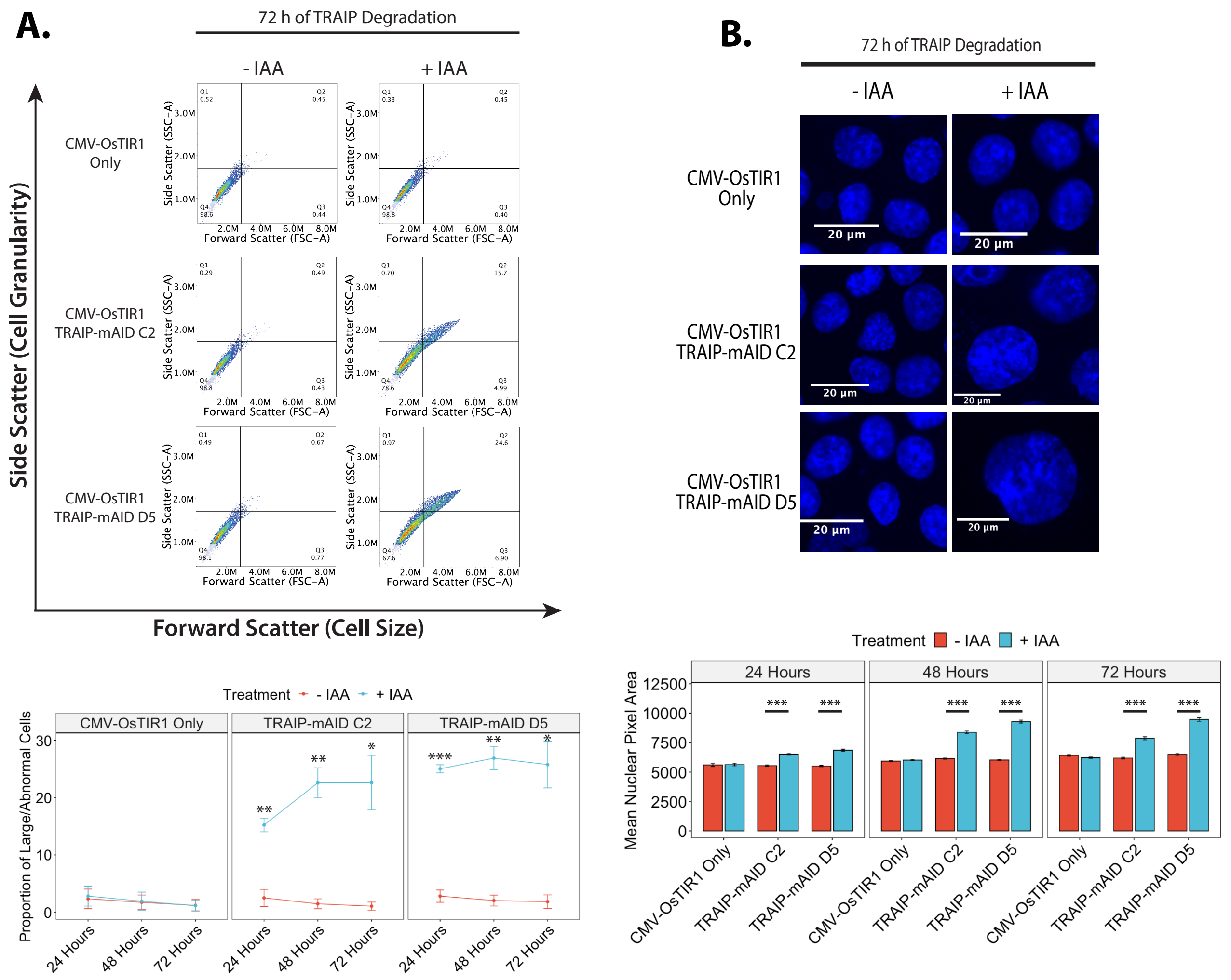


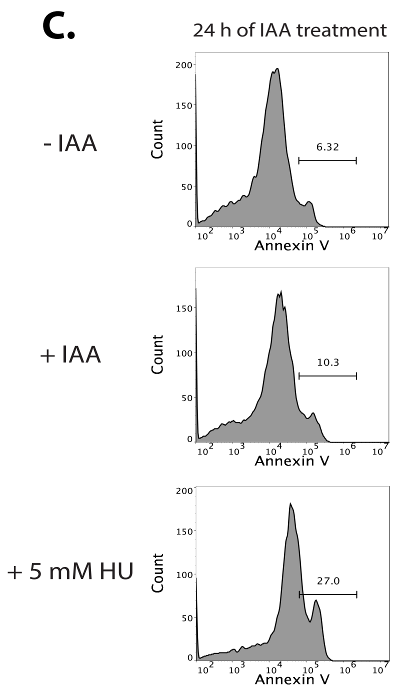


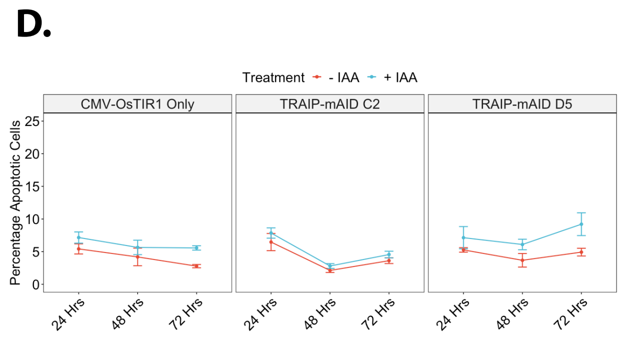


**Supplementary Figure 4. Analysis of consequences of TRAIP depletion. (A)** The size of TRAIP degron cells was analysed by flow cytometry after auxin addition. The example FACS plots after 72 h of IAA treatment (top) and quantification over 24, 46 and 72h TRAIP degradation n=3 (bottom) is presented. (B) The size of nuclei was analysed by DAPI staining and fluorescence microscopy. Example pictures after 72 h of IAA treatment (top) and quantification over 24, 46 and 72h TRAIP degradation n=3 (bottom) is presented. **(C)** Annexin V staining assay validation. TRAIP-mAID cell lines were treated optionally with auxin for 24 h or HU for 24 h. Annexin V positive cells were detected by FACS. Example FACS profiles presented.
**(D)** Quantification of percentage of TRAIP degron cells positive for Annexin V after 24, 48 or 72 h of auxin treatment.

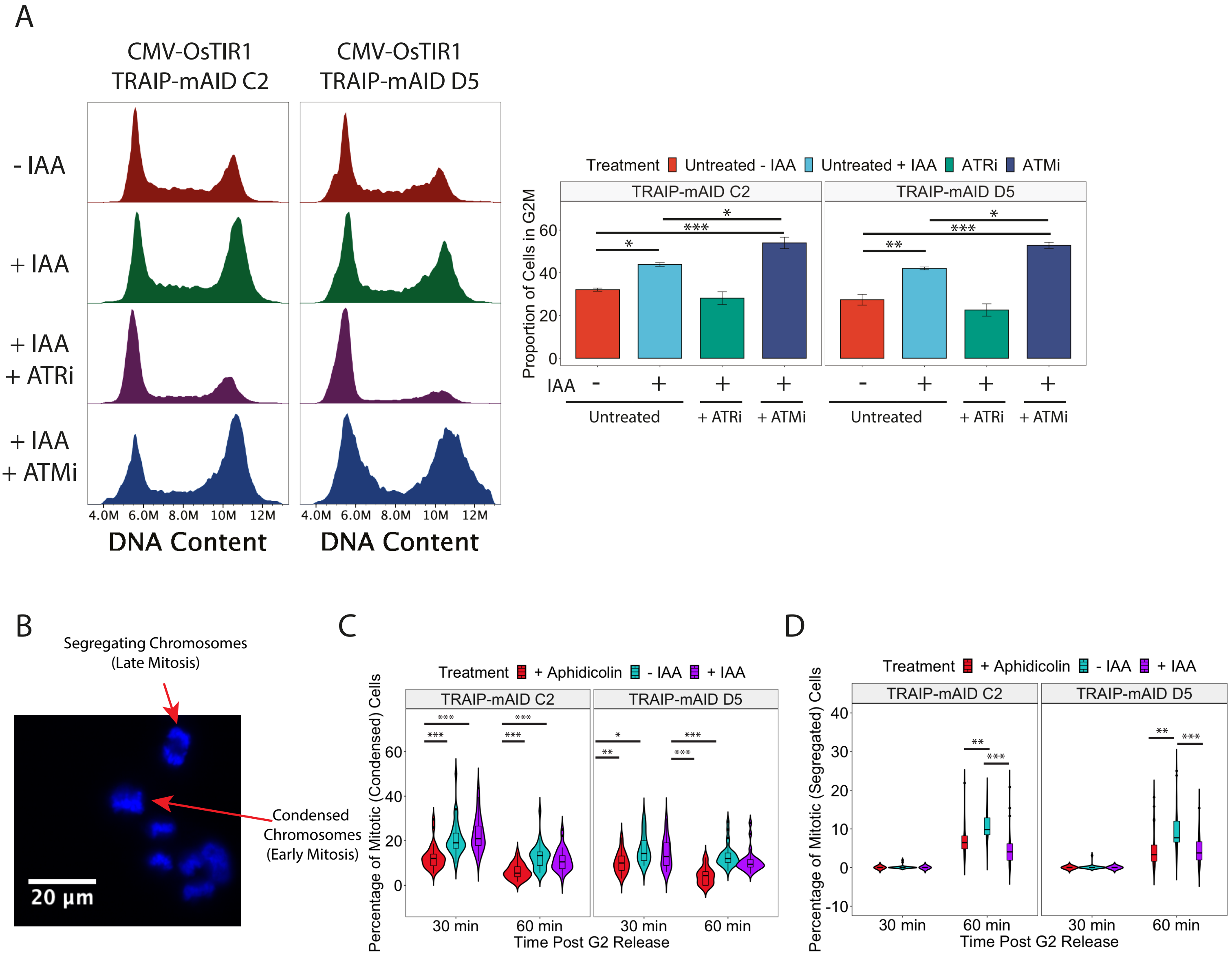


**Supplementary Figure 5. (A) G2 arrest in TRAIP depleted cells is dependent on functioning ATR checkpoint pathway.** TRAIP degron clones were treated for 24 h with auxin and optionally with ATRi or ATMi. Cell cycle profile of cells was analysed by FACS. Example of FACS plots (left) and quantification of cells in G2/M stage of the cell cycle n=3 (right).
**(B-D) TRAIP regulates mitotic progression.** Cells were arrested in G2 with RO-3306. TRAIP was degraded, cells released into mitosis and progression through mitosis observed at 30 and 60 min post release. As a control cells were treated with aphidicolin during RO-3306 treatment to slow down progression through the cell cycle. **(B)** example of detection of cells with condensed and segregating chromosomes. **(C)** Quantification of percentage of cells with condensed chromosomes n=3. **(D)** Quantification of percentage of cells with segregating chromosomes n=3.

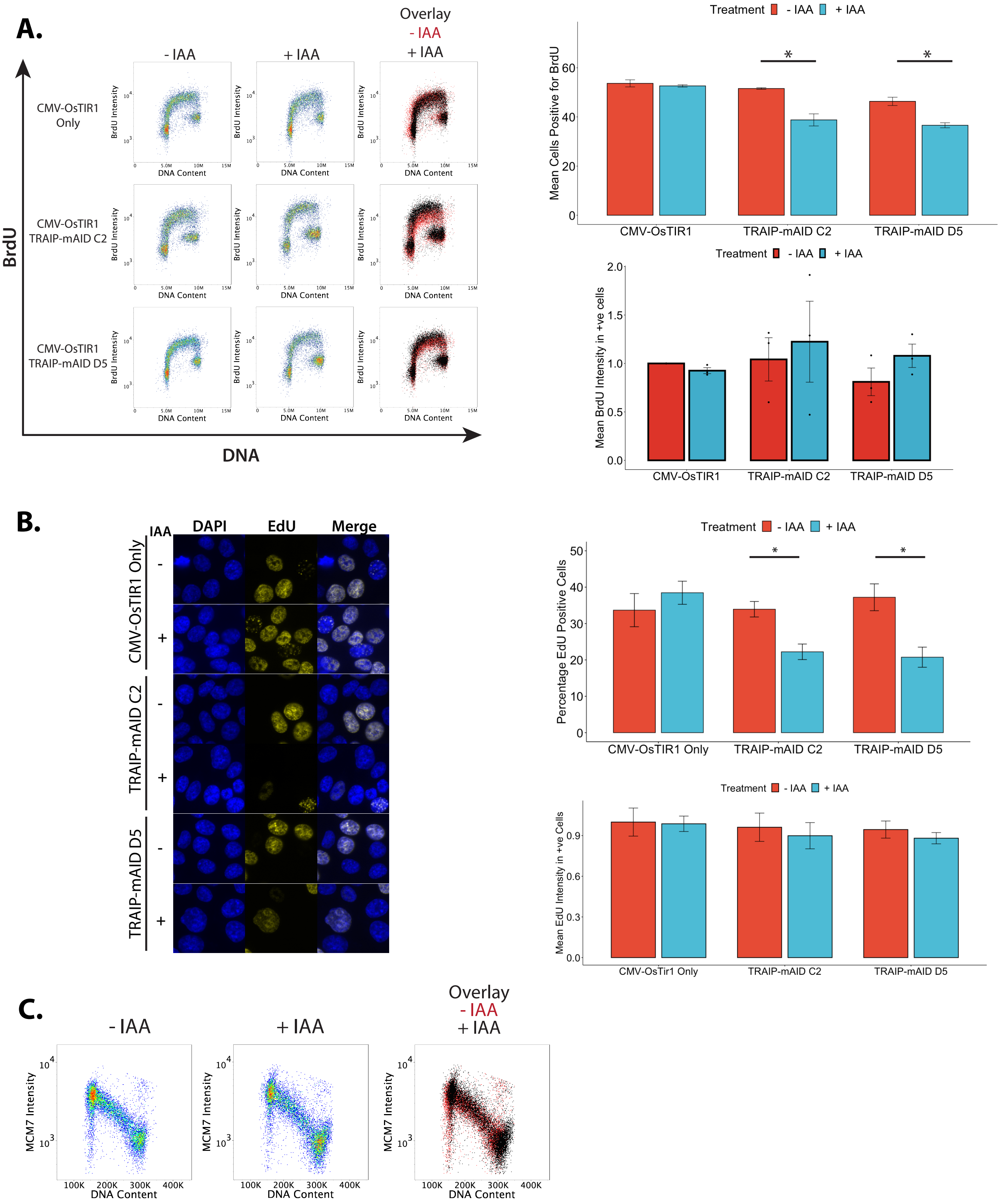


**Supplementary Figure 6. TRAIP degradation does not affect global DNA replication. (A)** Cells optionally treated with IAA for 24 h were pulsed with BrdU for 1 h and BrdU incorporation into DNA analysed by FACS. Example FACS plots are presented (left) and quantification of number of cells incorporating BrdU and the level of BrdU incorporation in replicating cells over n=3 (right). mean and SEM. Statistical analysis was carried out using t. tests, and any significant differences discovered is summarised on the plot (TRAIP-mAID C2: p = 0.0331; TRAIP-mAID D5: p = 0.0126). **(B)** Cells optionally treated with IAA for 24 h were pulsed with EdU for 1 h and EdU incorporation into DNA detected by immunofluorescence. Example microscopy images are presented (left) and quantification of number of cells incorporating EdU and level of EdU signal in replicating cells over n=3 (right). Statistical analyses was carried out on the data using t. tests and significant differences are summarised on the plots (TRAIP-mAID C2: p = 0.0178; TRAIP-mAID D5: p = 0.0264). **(C)** Cells were optionally treated with IAA for 24 h, nuclei extracted and the level of chromatin bound Mcm7 analysed by FACS.

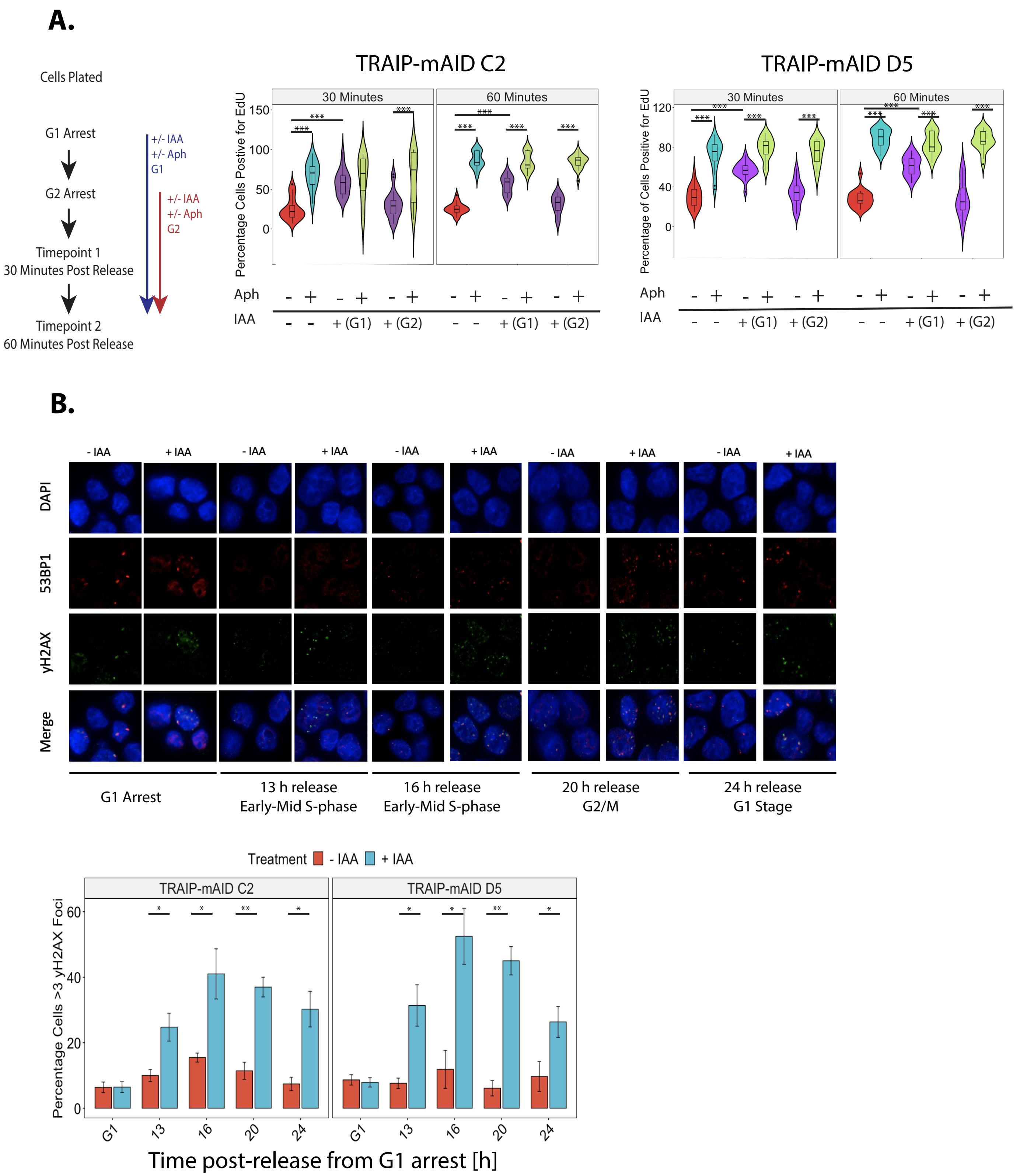


**Supplementary Figure 7. DNA replication defect and DNA damage observed in cells after TRAIP depletion. (A)** TRAIP Degradation Prolongs DNA Replication into G2/M. Cells were synchronised and treated as depicted in the schamatic (left). Quantification of the total proportion of cells positive for EdU incorporation for experiments using TRAIP-mAID C2 and TRAIP-mAID D5 (n = 3). Any differences between groups were determined using one-way ANOVA testing and post-hoc pairwise comparisons, indicated on the plots ($p > 0.001$ for all comparisons shown). **(B)** TRAIP was degraded just before S-phase as in (A). At indicated time points the DNA damage foci (γH2AX and 53BP1 foci) were analysed by immunofluorescence. Example images are presented (top) and quantification of percentage of cells with over 3 γH2AX foci over n=3 experiments (bottom). Significant differences (t.tests) are summarised on the graph for TRAIP-mAID C2 (13 Hrs post release: $p = 0.028$; 16 Hrs post release: $p = 0.037$; 20 Hrs post release, $p = 0.0016$; 24 Hrs post release, $p = 0.019$) and TRAIP-mAID D5 (13 Hrs post release: $p = 0.028$; 16 Hrs post release, $p = 0.010$; 20 Hrs post release, $p = 0.0019$; 20 Hrs post release, $p = 0.032$).

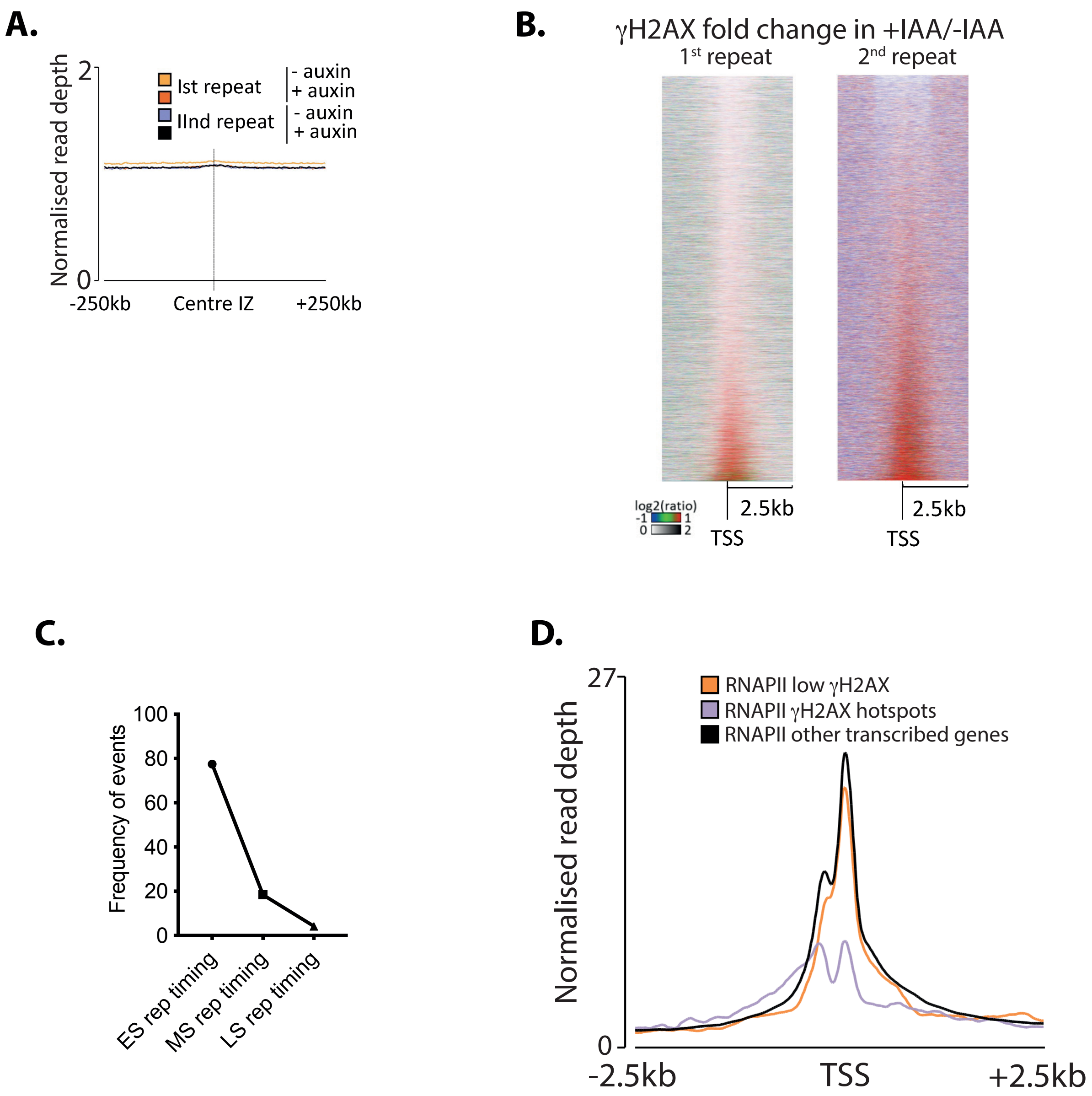


**Supplementary Figure 8. The increase of γH2AX upon TRAIP degradation does not correlate with replication features but with transcription start sites. (A)** Correlation of γH2AX signal with replication initiation and termination zones as determined by Daigaku et al. 2022. **(B)** Heatmaps of γH2AX signal centered on transcription start sites (TSS). TSS sorted by γH2AX fold change in +IAA/-IAA. Two independent repeats are presented. **(C)** Replication timing of hotspots. **(D)** RNAPII ChIP-seq peaks at TRAIP-depletion induced γH2AX hotspots suggest bidirectional DNA transcription from these TSS.

**A.**

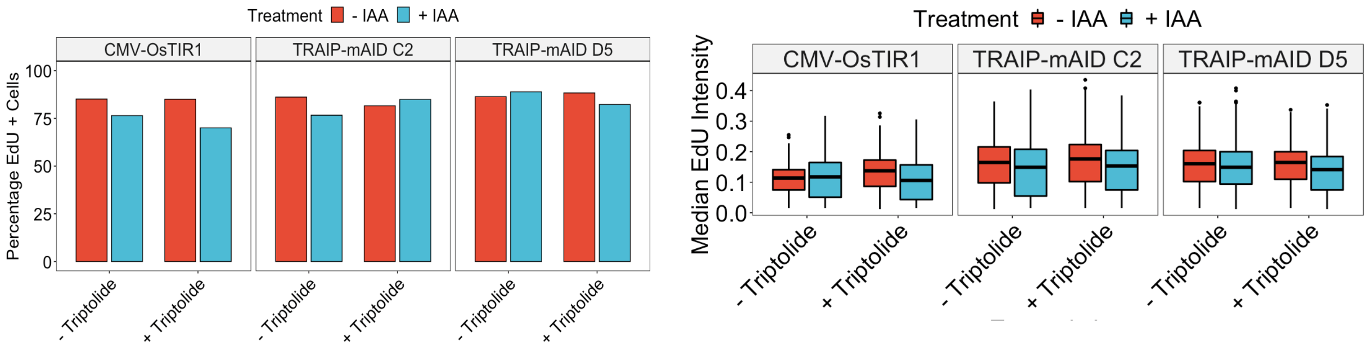


**B.**

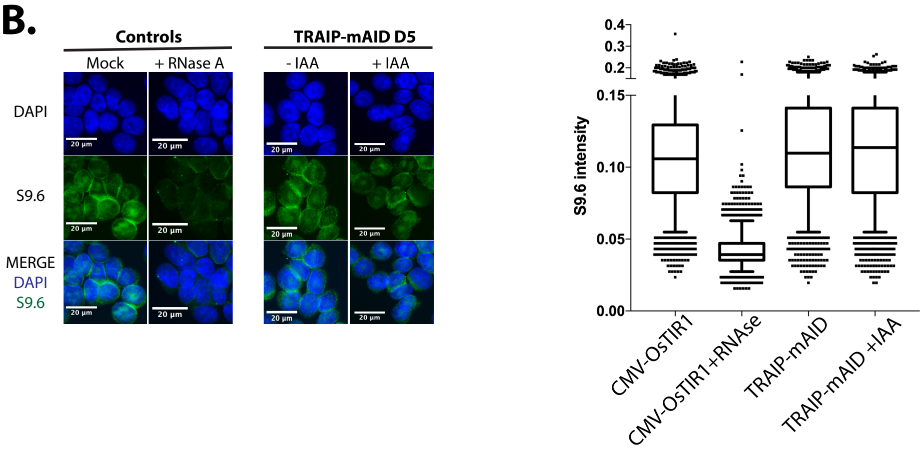


**Supplementary Figure 9. (A) Short Triptolide treatment in S-phase does not inhibit DNA replication.** Cells were synchronised in G1 with optional degradation of TRAIP for last 1 h of arrest and released into mitosis. Cells were then treated for 90 min with Triptolide, 3 h after release from G1 arrest. Cells were pulsed for 1 h with EdU and its incorporation was analysed using fluorescent microscopy. Shown are the overall percentage of cells positive for EdU (left), as well as median EdU intensity per cell (right). **(B) No changes in R-loops level was detected upon TRAIP depletion.** The S9.6 RNA:DNA hybrid antibody was used to detect any major changes to R-loop levels in the conditional degron cells. S-phase cells with and without TRAIP were stained for the S9.6 antibody and the respective fluorescence measured using Cell Profiler. Examples of immunofluorescence images (left). Shown is DAPI (Blue) and S9.6 (Green). Quantification of S9.6 intensity. Median value with 25% and 50% quartiles indicated by box whisker (n=3).

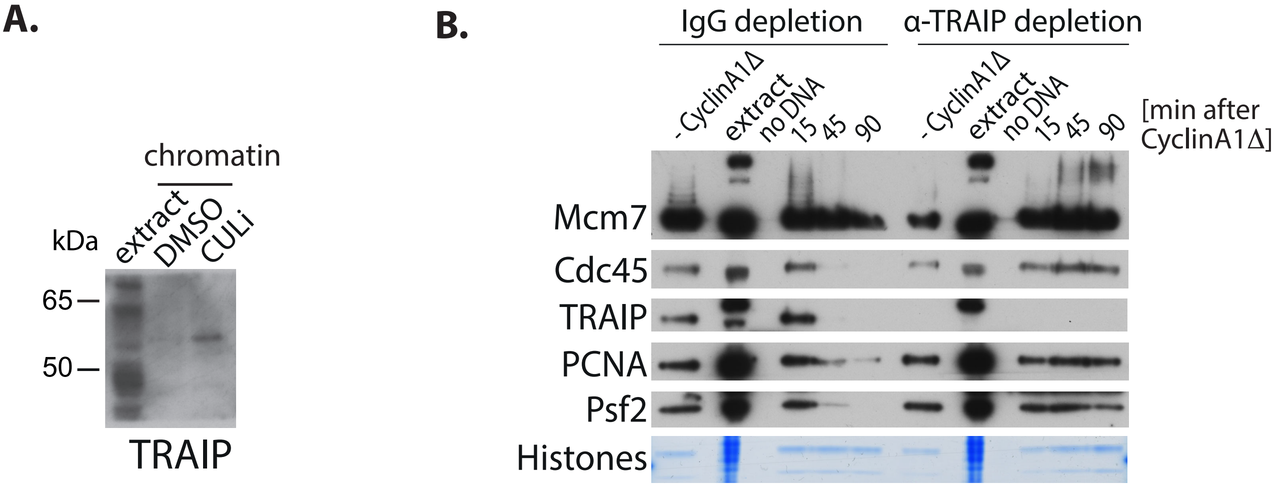


**Supplementary Figure 10. TRAIP depleted Xenopus egg extract is unable to unload replisomes in mitosis. (A)** Characterisation of *Xl*TRAIP antibody raised for this study. The level of TRAIP in the egg extract is low and all of our TRAIP antibodies recognise other bands in full egg extract. We have therefore also resolved samples of chromatin with accumulated replisomes (MLN4924 treatment) to confirm that the band between 65 and 50 kD is indeed behaving as expected from TRAIP. **(B)** TRAIP depleted extract cannot unload replisomes in mitosis. Non-specific IgG or α-TRAIP depleted egg extract was used to replicate DNA to completion in presence of neddylation inhibitor MLN4924, which inhibits activity of Cullin type ubiquitin ligases and blocks S-phase pathway of replisome disassembly. Once replication was completed, extracts were supplemented with Cyclin A1Δ to stimulate progression into mitosis. Chromatin was isolated at indicated timepoints and replisome presence on chromatin was monitored by western blotting with indicated antibodies. In mock depleted extract CMGs (as visualised by Cdc45, Psf2) are removed from chromatin, while in α-TRAIP depleted egg extract they are retained on chromatin due to lack of TRAIP.

**Supplementary Table 1.** Summary of the significant GOTerms identified from the genes isolated using γ-H2AX ChIP sequencing

| Analysis Type | Gene Ontology Term | Count (# Genes) | Frequency (%) | Corrected P-Value (Benjamini) |
|---|---|---|---|---|
| Biological Process (GOTerm BP) | positive regulation of transcription from RNA polymerase II promoter | 55 | 10.5 | 0.0014 |
| Biological Process (GOTerm BP) | negative regulation of transcription from RNA polymerase II promoter | 42 | 8 | 0.0077 |
| Biological Process (GOTerm BP) | ephrin receptor signaling pathway | 12 | 2.3 | 0.013 |
| Biological Process (GOTerm BP) | vascular endothelial growth factor receptor signaling pathway | 11 | 2.1 | 0.013 |
| Biological Process (GOTerm BP) | transcription from RNA polymerase II promoter | 31 | 5.9 | 0.034 |
| Biological Process (GOTerm BP) | transcription, DNA-templated | 81 | 15.4 | 0.045 |
| Biological Process (GOTerm BP) | regulation of actin cytoskeleton organization | 8 | 1.5 | 0.071 |
| Biological Process (GOTerm BP) | covalent chromatin modification | 12 | 2.3 | 0.071 |
| Biological Process (GOTerm BP) | protein phosphorylation | 27 | 5.1 | 0.08 |
| Molecular Function (GOTerm MF) | protein binding | 289 | 55 | 0.0026 |
| Cellular Component (GOTerm CC) | nucleus | 206 | 39.2 | 0.000000016 |
| Cellular Component (GOTerm CC) | nucleoplasm | 114 | 21.7 | 0.000047 |
| Cellular Component (GOTerm CC) | cytoplasm | 175 | 33.3 | 0.0041 |
| Cellular Component (GOTerm CC) | cytosol | 117 | 22.3 | 0.017 |
| Biological Process (GOTerm BP) | positive regulation of transcription from RNA polymerase II promoter | 55 | 10.5 | 0.0014 |
